# Co-translational O-glycosylation driven by GALNTs spatial reprogramming fosters pancreatic cancer growth

**DOI:** 10.64898/2026.09.04.749345

**Authors:** Rebecca Bennion, Eugenie Lohmann, Giacomo Del Rio, Lucile Rouyer, Sahar El Amrani, Malgorzata Kowalczewska, Xavier Le Guezennec, Eric Mas, Nelson Dusetti, Alex Chauvin, Richard Tomasini, Sophie Vasseur, Fabienne Guillaumond, Brice Chanez, Martin J. Humphries, Sergey Y. Vakhrushev, Frederic A. Bard

## Abstract

Signal-driven relocation of GalNAc-transferases (GALNTs) from the Golgi to the ER, termed GALA, promotes tumour growth, but its effects on glycosylation are unclear. Unlike N-glycosylation, which is co-translational, O-glycosylation initiates post-translationally in the Golgi. Here we show that GALA subverts this arrangement in pancreatic ductal adenocarcinomas (PDAC) and in murine pancreatic tumours, where it stimulates growth. Quantitative glycoproteomics on a cellular model reveals a substantial expansion of the O-glycoproteome, consisting in thousands of sites across hundreds of proteins, ER-resident proteins, cell-surface receptors, secreted factors, and extracellular matrix components. Profiling of murine tumours and patient-derived xenografts confirms widespread activation and cross-species conservation of glycosylation patterns. Structural analysis reveals that GALA-specific residues have solvent accessibility as low as N-glycosylation sites, and below Golgi O-glycosylation or phosphorylation sites, indicating that ER O-glycosylation occurs co-translationally. By inverting the normal temporal sequence of folding and glycosylation, cancer cells generate alternative glycoforms that foster tumour growth.

## INTRODUCTION

Pancreatic ductal adenocarcinoma (PDAC), the most common form of pancreatic cancer, is the third leading cause of cancer-related mortality, with a five-year survival rate of only 8–10%^1^. This poor prognosis reflects both late diagnosis and the limited efficacy of current therapies. A defining feature of PDAC is its complex tumour microenvironment (TME), characterised by extensive extracellular matrix (ECM) deposition and a heterogeneous mixture of tumour, stromal, and immune cells^2^. Interactions among these components play central roles in tumour progression and therapeutic resistance^3^. Tumor growth depends on coordinated cell–cell and cell–ECM interactions mediated by cell-surface and secreted proteins from cancer cells. These proteins are synthesised first in the endoplasmic reticulum (ER) before being modified by N- and/or O-glycans.

In mammalian cells, these two main glycosylation systems differ fundamentally in their compartmentation. N-glycosylation is initiated in the ER by co-translational addition of an oligosaccharide ^4^. By contrast, O-linked glycosylation is initiated in the Golgi by addition of a single N-acetylgalactosamine (GalNAc), forming the Tn glycan. The responsible enzymes, the family of GalNAc-transferases (GALNTs), encounter their substrates after folding and complex formation has completed.

O-linked glycosylation is frequently affected in cancer, with elevated Tn expression a hallmark of many carcinomas, first described more than four decades ago ^5^. The mechanisms underlying this increase and the identity of affected proteins remain debated ^6^. A classic explanation of elevated Tn is a defect in downstream glycan elongation, with the inability to extend Tn and form the core 1 disaccharide (Galβ1–3GalNAc, or T antigen)^7^. Downregulation of C1GALT enzyme or its chaperone has been proposed to drive high Tn in various carcinomas, including PDAC ^8^. This mechanism would leave the number of glycosylation sites in the proteome unchanged and eliminate extended O-glycans.

Alternatively, elevated Tn antigen levels can result from activation of the GALNT Activation (GALA) pathway, in which GALNTs relocate from the Golgi to the ER, leading to increased addition of the Tn glycan onto ER-resident proteins ^9,10^. Unlike loss of C1GALT1, which leaves the underlying proteome of glycosylated sites unchanged, GALA actively expands the pool of Tn-bearing proteins by exposing new ER-resident substrates to GALNTs. GALA is activated downstream of oncogenic signaling pathways, including Src, EGFR, and PDGFR ^9,11,12^.

Specifically, Src induces GALNT relocation via tubulo-vesicular structures formed by the retrograde trafficking factor GBF1, which is activated through tyrosine phosphorylation ^13^. GALA has been documented in liver and breast cancer. Experimental activation of GALA, through expression of an ER-targeted GALNT1 or GALNT2 enzyme, strongly promotes tumor growth and metastasis ^14,15^. Mechanistically, this spatial reprogramming of O-glycosylation markedly enhances ECM degradation. This degradative activity is mediated in part by hyperglycosylation of the cell-surface collagenase MMP14, where additional O-glycosylation sites result in enhanced protease activity ^15^. In addition, GALA induces glycosylation of the ER-resident chaperone calnexin (Cnx); this glycosylation drives relocalization of a fraction of Cnx to the cell surface, where it contributes to ECM degradation by reducing disulfide bonds in ECM proteins ^16^. These two examples illustrate that organellar redistribution of GALNTs can regulate the activity of multiple, functionally unrelated proteins through hyperglycosylation. To date, it remains unclear whether this pathway is activated in other cancers, such as PDAC, which proteins are affected by hyperglycosylation in those contexts. At a more fundamental level, it is unknown why relocation of GALNTs to the ER leads to an overall increase in Tn levels. How is the GALNT activity limited when in the Golgi?

Here, we show that GALA is widely activated in human and mouse PDAC. Using glyco-engineered mouse cell lines, we show that ER glycosylation drives pancreatic tumor expansion and that ER-localised GALNT activity induces widespread remodelling of the O-glycoproteome. Analysis of mouse pancreatic tumors and patient-derived xenografts (PDX) confirms the presence of GALA-dependent glycoforms in human tumors with glycopatterns conserved across species. GALA increases the glycosylation of hundreds of proteins, including key regulators of cell adhesion and signalling. Analysis of glycosites in the context of protein 3D structure reveals a large increase in buried residues with ER-specific sites indicating that ER-targeted GALNTs modify substrates before they fold. Together, these findings indicate reorganisation of O-glycosylation initiation induces markedly different glycoforms by inverting the temporal sequence of folding and glycosylation.

## Results

### High Tn is detected in CK19+ cells in a majority of PDAC samples

To assess the prevalence of altered O-glycosylation in PDAC, a tissue microarray comprising 36 human PDAC tumours and eight normal pancreatic samples was analysed. Sections were stained with biotinylated *Vicia villosa* lectin (VVL), which specifically recognises the Tn antigen, and detected using fluorescent streptavidin (Fig. S1a). Compared with normal pancreas, most PDAC cores exhibited markedly increased VVL staining (Fig. 1a). Quantification of VVL-positive area above threshold revealed that 65% of tumour cores showed at least a two-fold increase relative to controls (Fig. 1b, S1b).

**Fig. 1.**
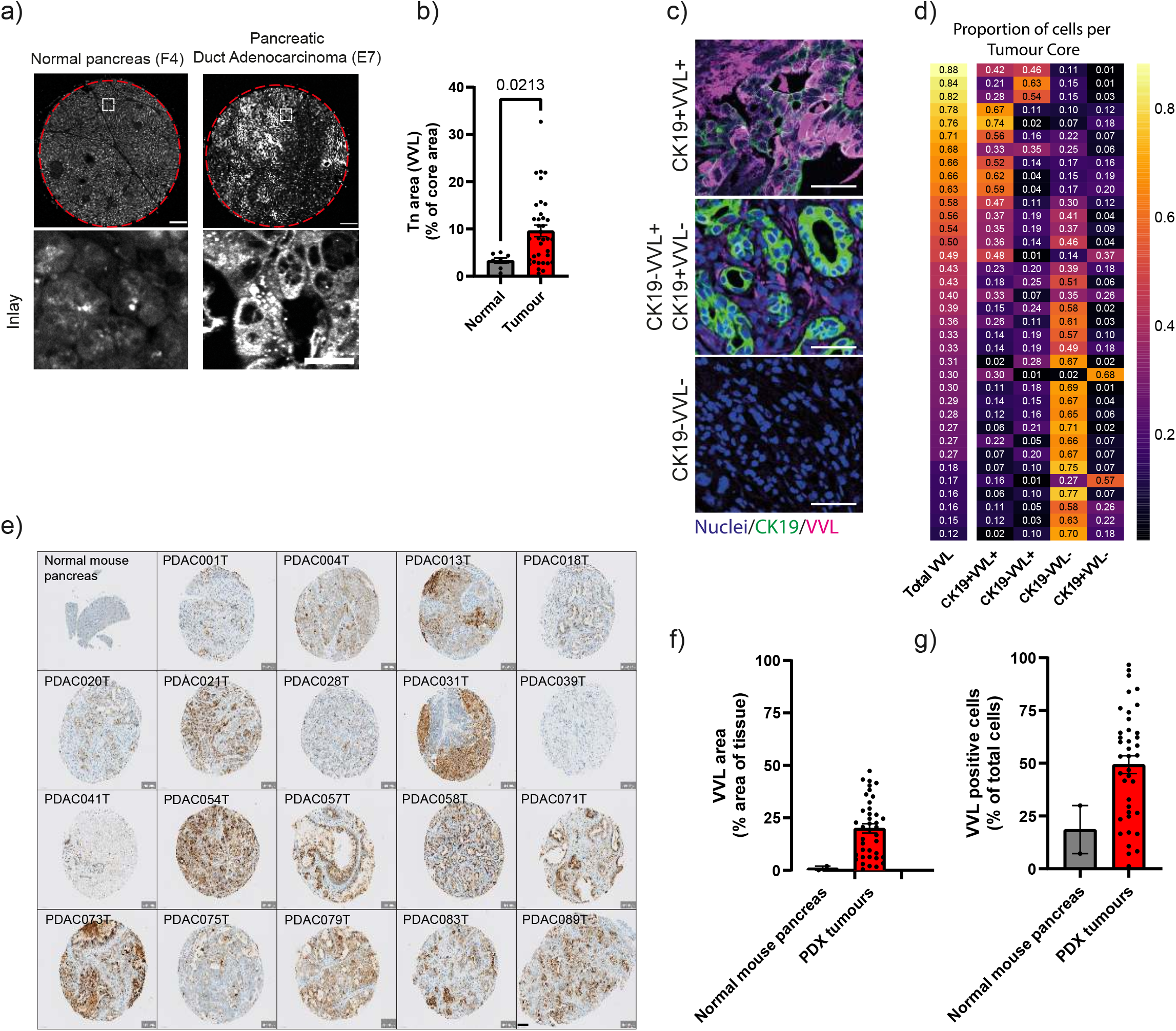
Tn glycan is elevated in human PDAC tissues. **a.** Representative images of a human tissue microarray (TMA) containing cores in duplicate from four normal and 18 pancreatic duct adenocarcinoma patients (upper panel; scale bar 200 µm). Inlay areas bottom panel (scale bar 20 µm). Tn glycan is stained with VVL. **b.** Quantification of Tn area in individual cores. Data points represent the area of VVL staining of the core as a percentage of the individual core area. Mean ± SEM is indicated. (p=0.0220; Normal vs tumour, Mann-Whitney test). **c.** Co-staining of VVL with the tumour epithelial marker, cytokeratin-19 (CK19). The TMA cores were classified into four categories based on co-staining: CK19+VVL+, CK19-VVL+, CK19–VVL– and CK19+VVL+. Nuclei are stained with Hoechst in blue, Tn in magenta and CK19 in green (scale bar 50 µm). **d.** Automated image analysis was applied to the TMA to derive the proportion of cells in each category. The heatmap is ordered with the proportion of cells positive for VVL (total VVL). Color code ranges from black 0 to yellow 1. **e.** 19 Patient-derived xenograft (PDX) samples from the PaCaOmics cohort alongside normal mouse pancreas were stained for the Tn glycan with VVL (scale bar 100 µm). **f.** The area of VVL staining per PDX tumour core was calculated as a percentage of the individual core area. Bar shown as mean ± SEM. Most PDX cores were in duplicate per sample, some duplicates were excluded in the analysis due to incomplete or damaged cores on the TMA. **g.** The percentage of VVL-positive cells was quantified per core. Bar shown as mean ± SEM.

To define the cellular origin of Tn expression, samples were co-stained for cytokeratin 19 (CK19), a ductal epithelial marker frequently overexpressed in PDAC. In many cores, elevated Tn expression was predominantly detected in CK19⁺ tumour cells, although CK19⁻/Tn⁺ stromal cells were also observed (Fig. 1c). A cell classification pipeline based on nuclear segmentation and marker intensity thresholding was developed to assign cells into four populations defined by CK19 and Tn status, enabling quantification of the heterogeneity observed visually in the cores (Fig. 1d, S1c,d). Across the cohort, 14 of 36 tumours contained at least 50% Tn⁺ cells, and 30 of 36 contained at least 20%. In most tumours, the majority of Tn⁺ cells were CK19⁺, with only three cases showing more than 30% CK19⁻/Tn⁺ cells. Conversely, most CK19⁺ cells were Tn⁺, although substantial intratumoural heterogeneity was observed, with several cores containing mixed CK19⁺/Tn⁺ and CK19⁺/Tn⁻ populations. A similar heterogeneous pattern was observed in pancreatic tumours from KPC mice, indicating conservation across species (Fig. S1e). We next examined Tn expression in 19 PDAC PDX from the PaCaOmics cohort using VVL staining ²⁹. Normal mouse pancreas used as control showed negligible staining, whereas PDX tumours displayed heterogeneous Tn expression, with most cores containing subsets of strongly Tn⁺ cells (Fig. 1e). Quantification of Tn⁺ area in PDXs yielded values comparable to those observed in the human PDAC tissue microarray (Fig. 1f,g). Overall, a high prevalence of increased Tn was observed, suggestive of GALA activation in most PDAC samples.

### PDAC cells display an ER pattern for Tn and GALNTs

To explore whether the increase in Tn levels was due to GALA, the TMA was co-stained with fluorescently labelled Helix Pomatia Lectin (HPL), another lectin with specificity for Tn, and the ER marker GRP78. Colocalisation was observed for both markers in high Tn cells, indicating that Tn is enriched in the ER (Fig. 2a). Next, the PDAC tumour cores were stained with a GALNT2 antibody and the Golgi marker GOLPH2 or, in adjacent sections, the ER marker GRP78 (Fig. 2b,c). While in some cells GALNT2 had the expected localisation at the Golgi, in other cells the enzyme had a clear ER pattern, indicating that GALNT2 relocalisation occurred (Fig. 2b,c). To note, past studies have shown that GALNT1 and 2 usually relocate indistinctively ^15^.

**Figure 2.**
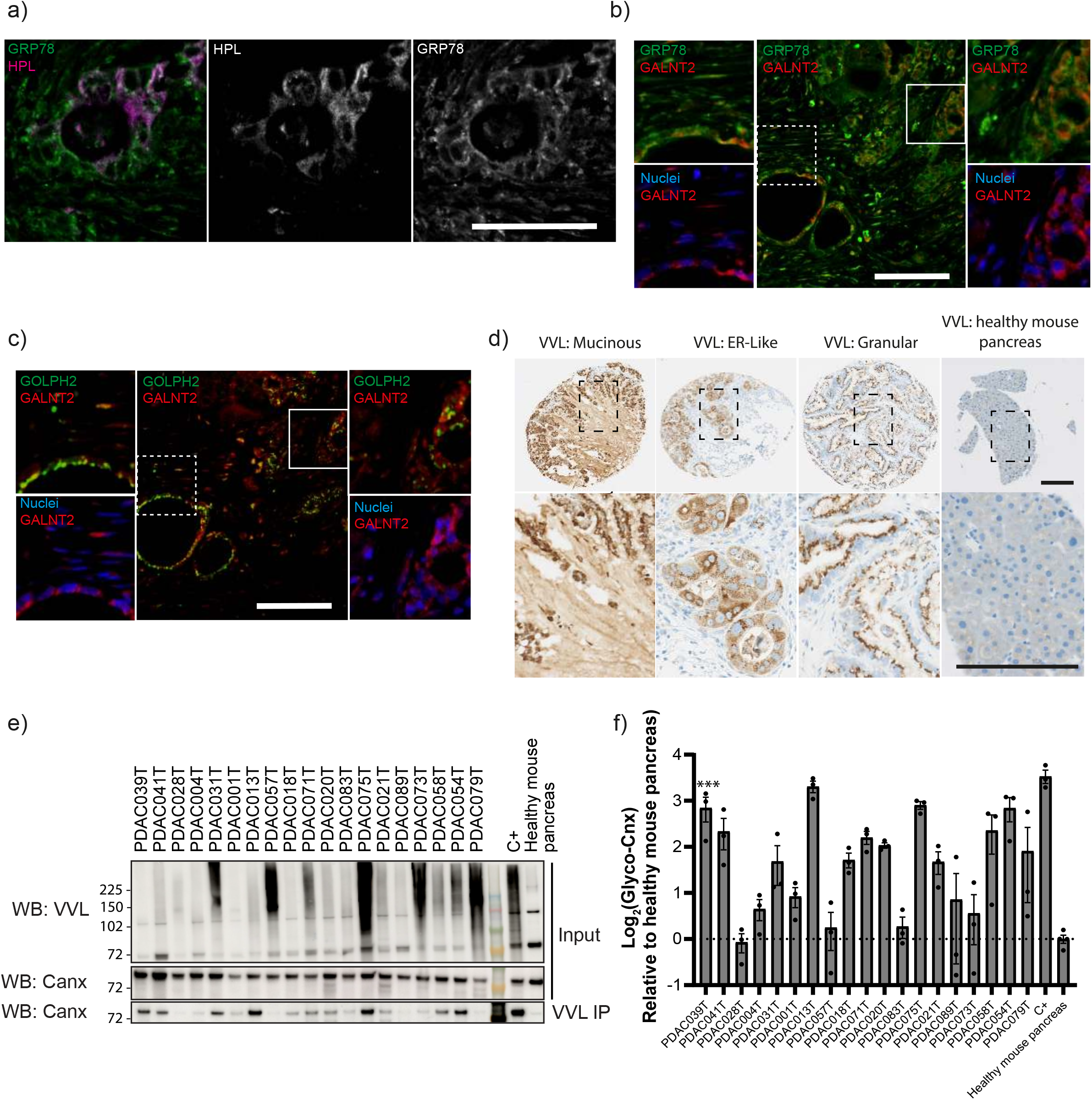
ER-localised expression of Tn glycan and glycosylated Calnexin in PDAC tumours and PDX models. **a.** PDAC tissue stained with Tn glycan (HPL) and the ER marker GRP78 (Scale bar 50 µm). **b** GALNT2 (red) was co-stained with the ER-marker GRP78 (green) in PDAC patient tumours (scale bar 50 µm). Dashed inlay (left panel) indicates a Golgi-localised GALNT2 pattern, whereas solid inlay (right panel) indicates an ER-localised GALNT2 pattern. Nuclei stained with Hoechst. **c.** Using an adjacent serial section from **b.** GALNT2 (red) was co-stained with the Golgi marker GOLPH2 (green). Dashed inlay (left panel) indicates the Golgi-localised GALNT2 and the solid inlay (right panel) indicates the ER-localised GALNT2. **d.** Immunohistochemistry VVL staining in healthy mouse pancreas and PDX PDAC tumours. Three main VVL staining patterns are shown: mucinous, ER-like and granular. A zoomed inlay is shown (right panel; scale bar 100 µm). **e.** Immunoblot analysis of the levels of Tn-modified Calnexin (glyco-canx) in PDX PDAC samples, normal mouse pancreas and the high GALA control SUM-190 cell-derived xenograft (CDX) tumour (C+). Lysate (input) and elution of VVL-agarose enrichment (VVL IP) are shown. Immunoblot (WB) was performed for VVL and Calnexin (Canx). **f.** Quantification of the glycosylated Calnexin (Glyco-Canx) VVL IP analysis: n = 3 for each PDX, n = 4 for healthy mouse pancreas, and n = 3 for MDA-MB-231 CDX (C+). Values are presented as Mean Log₂(Glyco-Canx) ± SEM, normalized to healthy mouse pancreas. Statistical significance was assessed using Ordinary one-way ANOVA with Dunnett’s multiple comparisons test (vs. healthy mouse pancreas): **** p < 0.0001, *** p = 0.0004–0.0001.

A similar ER pattern of Tn was observed in many of the PDX cores, although some Tn staining also appeared concentrated in granular structures and was also found extracellularly (Fig. 2d). To measure GALNT activity in the ER with a semi-quantitative approach, the GALA-dependent glycosylation of the ER-resident protein Calnexin was exploited^16^. Using lysates of frozen samples of the 19 PDX, a pulldown of Tn-bearing proteins using VVL was performed followed by Calnexin immunoblotting (Fig. 2e, S2b). This approach demonstrated that 12/19 cores had an average 1.5 log_2_-fold (approximately 2.8-fold) increase in glyco-Calnexin over healthy mouse pancreas (Fig. 2f). Together, these results strongly suggest that a major driver of high Tn expression in PDAC is the relocation of GALNTs to the ER.

Alternative mechanisms for high Tn levels have been suggested, such as the transcriptional downregulation of C1GALT1 or its chaperone, C1GALT1C1 (also known as Cosmc), which are responsible for the modification of Tn and synthesis of the core-1 disaccharide. Transcriptomic analysis of these enzymes in the PDX did not reveal any significant variations across samples (Fig. S2a). Several GALNTs were expressed consistently and at similar levels as C1GALT1, in particular GALNT 1-7 and 10-12 (Fig. S2a). Other GALNTs, such as GALNT8, 9, 13, 14 and 18 had lower and more variable expressions.

### Expressing an ER-targeted GALNT1 in KPC47 cells results in high Tn levels

To model GALNT relocalisation, an ER-targeted GALNT1 construct (ER-G1) generated by fusing GALNT1 to the C-terminal region of CD74 (p33) was used ^14^. Control constructs included GFP, wild-type GALNT1 (WT-G1), and a catalytically inactive ER-targeted mutant (ER-G1 H211D) (Fig. 3a). GALNT constructs were placed under the control of a tetracycline-responsive enhancer to enable inducible expression, whereas GFP was expressed constitutively. Stable cell lines were generated in the pancreatic cancer mouse cell line KPC47, which has low endogenous GALA. Doxycycline induction resulted in comparable expression levels of the three GALNT1 derived proteins (Fig. S3a). Immunofluorescence analysis confirmed ER localisation of ER-G1, whereas WT-G1 remained restricted to the Golgi apparatus (Fig. 3b). Expression of ER-G1 was sufficient to induce a marked elevation of Tn levels, whereas the catalytically inactive ER-G1 H211D failed to do so. WT-G1 expression resulted in only a modest increase in Tn (Fig. 3b).

**Figure. 3.**
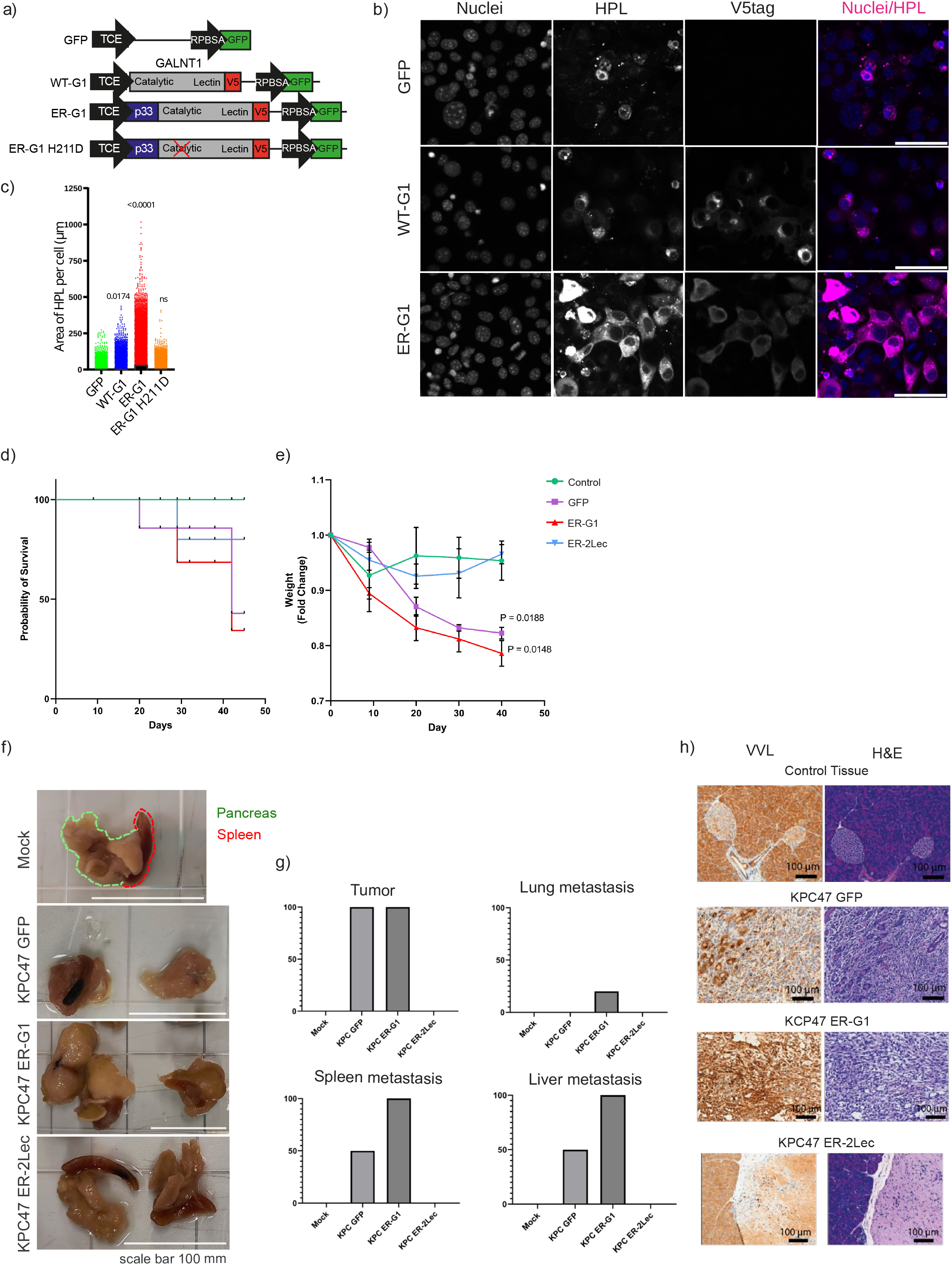
ER-targeted GALNT1 induces Tn glycan expression in pancreatic cancer cells. **a.** Schematic of the Sleeping Beauty gene expression constructs used in this study. Tetracycline-responsive promoter (TCE) controls empty vector, wild-type GALNT1 (WT-G1), ER-localised GALNT1 (ER-G1) tagged to the ER with the p33 domain of MHC class II protein and ER-localised catalytic mutant (ER-G1 H211D) ER-G1, all constructs are tagged with V5. The GFP is under control of the synthetic promoter RPBSA. **b.** The constructs were expressed in KPC47 cells, stained for Tn glycan (HPL) and V5 tag. Nuclei stained with Hoechst (blue) and HPL (magenta). Scale bar 50 µm. **c.** Quantification of HPL labels per cell in stably transfected KPC47 cells. Data points are individual cells. P-values are shown from ordinary one-way ANOVA with Dunnett’s multiple comparison test to GFP, n=3. **d.** Graph showing the probability of survival of mice after orthotopic injection and monitoring for 6 weeks. **e.** showing the mean weight loss ± SEM in fold change from weight pre-implantation at day 0 until day 42 of harvest. n=4-6 mice. Mantel-Cox log-rank test comparing the survival probability was performed (p=0.3063). Mixed-effects model analysis was performed to compare weight loss compared to control. **f.** Images show the pancreas with spleen attached, harvested from mice 6 weeks after implantation of KPC47 cells. Mock shows a pancreas after the implantation of Matrigel only. Scale bar 100 mm. **g.** Quantifications of tumour incidence (%), lung metastasis incidence (%), spleen metastasis (%) and liver metastasis (%). n=4-6 mice. **h.** Vicia villosa lectin (VVL) and H&E immunohistochemistry were performed on orthotopically injected pancreas. Images show the whole pancreas (Scale 5 mm) with inlays below (1 mm and 100 μm scale bars) for Mock, KPC GFP, KPC ER-G1 and KPC ER-2Lec 6 weeks after orthotopic implantations. n=4-6 mice.

Quantification of HPL-positive area per cell revealed that ER-G1 expression induced approximately eight-fold increase in Tn in cells, compared with a two-fold increase upon WT-G1 expression (Fig. 3c). Consistent with activation of the GALA pathway, ER-G1 expression also increased levels of Tn-modified Calnexin (Fig. S3b). Together, these results demonstrate that ER localisation of GALNT1 is sufficient to drive high Tn expression and highlight subcellular localisation of GALNTs as a key determinant of O-glycosylation changes in PDAC-derived cells.

### GALA O-glycosylation is essential for orthotopic tumor growth

In order to assess the impact of GALA on pancreatic tumor growth, an orthotopic implantation mouse model was used. KPC47 expressing GFP, ER-G1 or ER-2Lec were implanted into the pancreas of C57BL/6 with 10,000 cells in matrigel. ER-2Lec is an ER-specific inhibitor of O-glycosylation derived from the lectin domain of GALNT2 ^14^. Mice were administered doxycycline throughout the experiment and survivors were euthanised 6 weeks post-implantation. KPC ER-G1 and GFP had a survival probability of less than 50% and suffered significant weight loss compared to matrigel only injected mice (control) (Fig. 3d,e). ER-G1 expression resulted in larger tumors than GFP and increased metastasis to the spleen, liver and lung (Fig 3.f,g). Histologically, GFP-KPC cells produced tumors with prominent, irregular glands, with moderate differentiation. By contrast, ER-G1 cells produced very few gland structures, with a more sheet-like pattern.

KPC ER-2Lec implanted mice did not develop any tumor and did not show any weight loss (Fig. 3f,g). Histological analysis revealed the implanted cells had remained in the matrigel and not invaded the surrounding tissue. To note, ER-2Lec expression does not impair cell proliferation in vitro in a measurable manner, indicating that tumor growth inhibition is non cell autonomous.

### ER localisation of GALNT1 extensively expands the repertoire of O-glycoproteins

To assess how ER localisation of GALNT1 alters the glycoproteome, we compared KPC47 cells expressing ER-G1, WT-G1 or GFP. Following neuraminidase treatment and trypsinisation, glycopeptides were enriched using jacalin, which captures both Tn- and T-modified peptides (Fig. 4a). Quantitative glycoproteomics was performed by data-independent acquisition (glyco-DIA) workflow, informed by data-dependent acquisition (DDA)-derived libraries from jacalin-enriched ER-G1 lysates (6,458 precursors, 3,416 peptide sequences, 478 glycoproteins), enabling sensitive mapping of the O-glycoproteome (Fig. 4a, Supplementary Table 1) ^17^. Proteins/glycopeptides identified by glyco-DIA were filtered for presence in ≥3/5 biological replicates. Proteomic analysis identified a comparable number of proteins across all cell lines (∼5,000 each; Fig. 4b, Supplementary Table 1), with reproducibility (coefficient of variation (CV) 0.92–0.99) and comparable abundance across lines (Fig. S4a). By contrast, glycopeptide analysis revealed a 3.6-fold increase in KPC47-ER-G1 versus KPC47-GFP (600 to 2,200 on average; Fig. 4b), reproducible across replicates (Fig. S4b). KPC47-WT-G1 was more similar to GFP than to ER-G1, showing only a limited increase in glycopeptides abundance.

**Figure 4.**
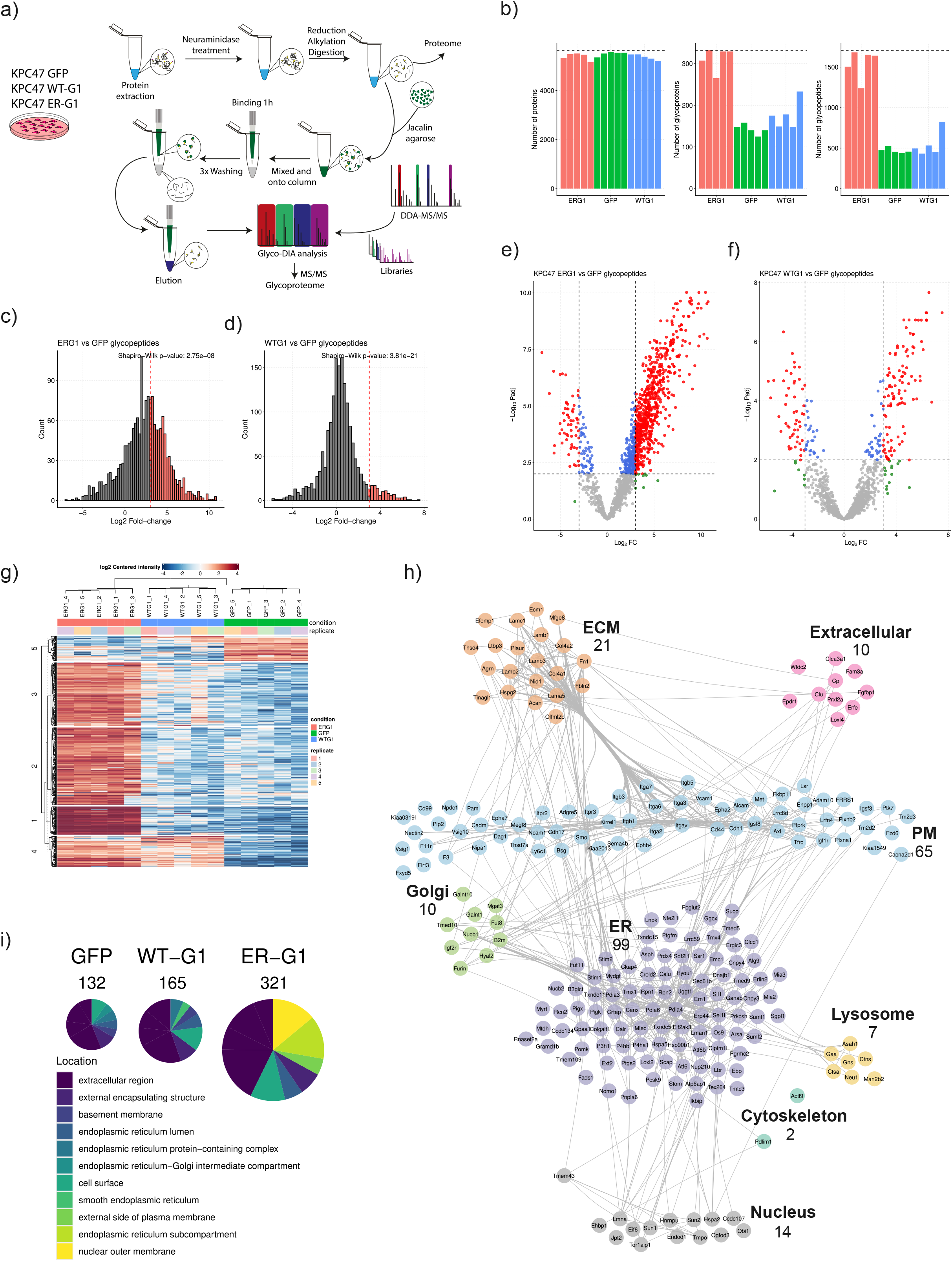
**a**. Schematic of the short lectin weak affinity (sLWAC) protocol coupled with glyco data-independent acquisition (glyco-DIA). Data-dependent acquisition (DDA) libraries from KPC47 cells were acquired prior. The proteome and glycoproteome of KPC47 GFP, WT-G1 and ER-G1 and 18 PDX PDAC tumours were analysed after enrichment with Jacalin agarose. **b.** Number of proteins, glycopeptides and glycoproteins detected. **c,d.** Glycopeptide-level histograms plots of Log_2_ fold-change between ER-G1 vs GFP and WT-G1 and GFP. Red dashed line is Log_2_ fold increase of 2. **e,f.** Glycopeptide-level volcano plot comparing ER-G1 vs GFP and WT-G1 and GFP. Differential analysis and a two-sided t-test was performed and the Benjamini-Hochberg false discovery rate (FDR) was calculated for the adjusted p-value. Log_2_ fold-change ≤ -2 and ≥ 2, and p-adjusted ≤ 0.05 represented by dashed lines, top-up15 proteins are annotated. **g.** Glycopeptide-level heatmap showing log_2_ centered intensity for differentially regulated glycopeptides (1% FDR). K-means clustering was performed on rows and Euclidean distance calculated between samples. **h.** Gene ontology biological processes analysis of each glycoprotein-level heatmap cluster. False discovery rate (p.adjust) is represented as a colour gradient from and dot size represents the fold enrichment. **i.** STRING PPI network of all KPC-derived candidate proteins (*n* = 240, *Mus musculus*), organized by subcellular compartment. Edges reflect STRING interaction confidence (score ≥ 0.4). Layout: Attribute Circle with edge bundling. **j** Pie charts showing the proportion of genes associated with the top enriched Gene Ontology cellular compartment terms for GFP, WT-G1 and ER-G1.

Mapping glycopeptides to proteins resulted in 330 glycoproteins on average in ER-G1 versus 150 in GFP (Fig. 4b, 4c, Supplementary Table 2). Overall, glycopeptide data were highly reproducible across replicates, yielding clustering in PCA (Fig. S4c,d). Differential analysis (log2 fold-change frequency) showed glycopeptide distributions were highly skewed for ER-G1 vs. GFP and ER-G1 vs. WT-G1. Almost half of glycopeptides were enriched with a >3 fold change (FC) in ER-G1 (Fig. 4c). By contrast WT-G1 vs. GFP, while still skewed, was centered on 0 FC (Fig. 4d). Volcano plots confirmed strong statistical significance for ER-G1 (Fig. 4e, f). Clustering of glycopeptides with p-adj ≤0.01 revealed two clusters specific for ER-G1 (1 and 2), while cluster 3 was upregulated in both WT-G1 and ER-G1; cluster 4 was downregulated in both, corresponding largely to T glycan-modified peptides (43% of all T-containing peptides identified) (Fig. 4g). The two ER-G1 clusters were enriched in proteins involved in integrin-mediated signaling (Fig. 4h). Glycopeptide modification analysis showed KPC47 cells mainly produce Tn (HexNAc) structures, with some T (Hex(1)HexNAc(1); Fig. S4e). In GFP cells, T and poly-T glycopeptides made up 23.4% of glycopeptides, versus only 8.8% in ER-G1; conversely, ER-G1 showed increased Tn and poly-Tn glycopeptides (Fig. S4e). These differences were not due to GALNT levels: GALNT1 abundance was similar in ER-G1 and WT-G1, and no other GALNT was induced by either construct (Fig. S4f). In sum, GALNT1 ER-localisation drives a marked increase in glycopeptide abundance, while GALNT1 overexpression alone has a limited effect.

### GALA impacts proteins in multiple cellular compartments, including key adhesion regulators and tumor promoters

To identify the protein classes affected by GALA, we performed over-representation analysis using Gene Ontology (GO) cellular compartment (CC) terms (Fig. 4i, S4g, Supplementary Table 3). In control cells, the most enriched terms were extracellular and cell surface, whereas ER-G1 expression enriched ER membrane and nuclear membrane terms, indicating a marked shift in the subcellular distribution of affected proteins. GO biological process (BP) analysis highlighted enrichment of terms associated with cell adhesion, protein maturation, and the ER-associated degradation (ERAD) pathway (Fig.4h, S4h). While ER-resident proteins were preponderant, they are not the only targets of ER-G1 (Fig. 4i). In terms of GO biological process, signaling receptor binding and cell adhesion emerged, while KEGG pathway analysis revealed integrin signaling and protein processing in ER (Fig S4i,j, Supplementary Table 4,5,6).

To better visualize the main glycoforms generated by GALA, 240 glycoproteins with glycopeptides significantly affected (log_2_ fold-change ≥ 2, p-adjusted ≤ 0.05) by ER-G1 expression were assigned a specific GO CC with manual curation when needed (Fig. S4k). The selected proteins were analysed for protein-protein interaction using the STRING database, revealing a highly interconnected network dominated by ER, cell surface and ECM proteins (Fig. 4j). ECM core basement membrane proteins are particularly enriched in ECM targets, with collagen IV, laminins, nidogen 1 (Nid1), perlecan (Hspg2), spondin-2 (Spon2), Tubulointerstitial nephritis antigen-like 1 (Tinagl1) and Extracellular matrix protein 1 (Ecm1). At the cell surface, transmembrane kinase receptors (Epha2, Axl, Met, Igf1r, Tgfbr1), GPCR (Smo), the ECM adhesion integrins, cell-cell adhesion molecules such as cadherins (CDH17 and CDH1), Vcam1, NCam1, Cadm1 and Nectin2 were found, suggesting a broad remodelling of surface activities (Fig. 4j, Supplementary Table 4,5,6). Most of these cell surface proteins have been involved in tumorigenic processes ^18–20^.

Collectively, these results demonstrate that ER-localised GALNT1 induces widespread remodelling of O-glycosylation across ER, cell surface, and ECM proteins, with particularly widespread targeting of cell-adhesion systems (Fig. S4l).

### GALA-type glycosylation in KIC tumors

Frozen PDAC from the spontaneous PDAC mouse model LSL-KrasG12D/+; Ink4a/Arffl/fl; Pdx1-Cre (KIC) at 9 weeks and age-matched pancreas from healthy mice were harvested and processed for glycoproteomics analysis as for KPC47 cell lines (Fig. 4a). Data derived from four healthy pancreas were highly similar and generated about twice as many glycopeptides than the five tumors (Fig. 5a). The data was highly differentiated between tumor and healthy and highly similar within the groups (Fig. 5b). Tumor specific glycopeptides were enriched in ER-resident proteins, while the healthy pancreas cluster was enriched in ECM and cell surface proteins (Fig. 5b, Supplementary Table7). Several ER-resident proteins found in the ER-G1 KPC47 dataset were enriched in KIC tumors, indicating activation of the GALA pathway (Fig. 5c).

**Figure 5.**
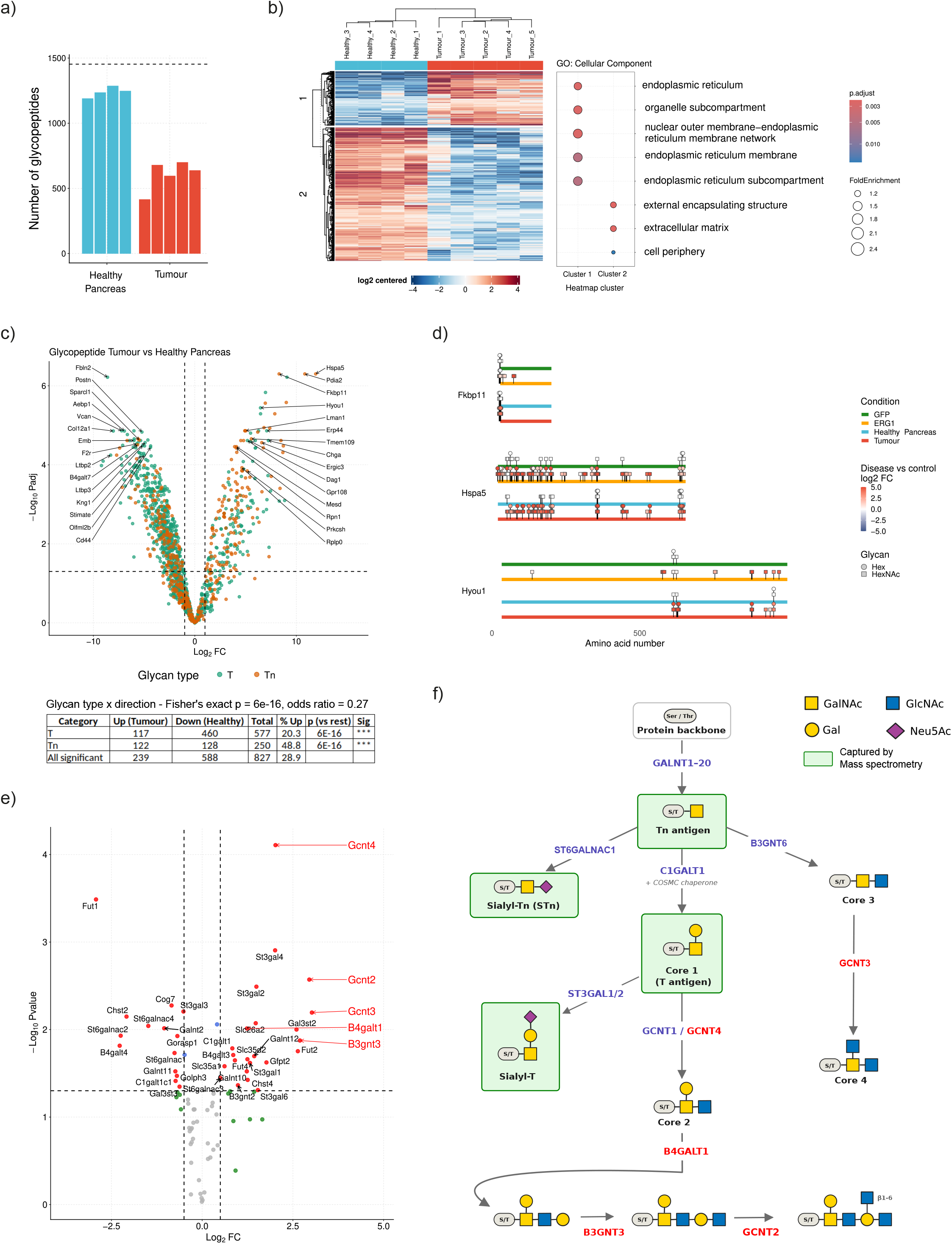
KIC tumors display evidence of GALA and core-2 extension. **a.** Number of glycopeptides per sample for each condition of Healthy Pancreas and Tumor. **b.** Glycopeptide-level heatmap showing log_2_ centered intensity for differentially regulated glycopeptides (1% FDR). K-means clustering was performed on rows and Euclidean distance calculated between samples. Gene ontology biological processes analysis of each glycoprotein-level heatmap cluster. False discovery rate (p.adjust) is represented as a colour gradient from and dot size represents the fold enrichment. **c.** Glycopeptide-level volcano plot comparing Healthy Pancreas to Tumour, peptides are coloured by glycan modification T(HexHexNac) or Tn(HexNac). **d.** Linear glycosite maps of top-up 5 Tumour-regulated glycoproteins (log₂FC ≥ 2, lowest adjusted p-value). Each protein displays two tracks (Healthy Pancreas and Tumor, top to bottom), positioned according to UniProt sequence annotations. Circle/Squares denote detected glycopeptide sites. UniProt domains are color-coded and labeled. **e.** Volcano plot of expression of O-glycosylation pathways genes. Highlighted in red are genes involved in extension of core 1 glycan. **f.** Schematic of O-glycosylation pathway with detectable and undetectable O-glycans

In contrast, ECM protein glycosylation diverged markedly from the KPC47 dataset. To investigate this discrepancy, we examined the relative distribution of Tn and T glycans. ER-resident proteins were predominantly modified with Tn glycan, whereas ECM proteins in healthy pancreas presented majoritarily T glycans (Fig 5c). Comparison of site-specific glycosylation patterns across three ER-resident proteins in KPC47 and KIC/HMP models revealed conserved glycosylation profiles (Fig. 5d).

We hypothesised that in KIC tumor cells, ECM and surface proteins were modified beyond the T glycan while transiting through the Golgi. The expression of O-glycosylation pathway enzymes was analysed systematically using previously acquired transcriptomic datasets from KIC-derived PDAC ^21^. Strikingly, several enzymes involved in the extension of O-glycans were over-expressed in KIC tumors compared to healthy pancreas: in particular GCNT2,3 and 4, and B4GALT1 and B3GNT3 (Fig. 5e). These enzymes extend the T glycan (core 1) into core 2 and further extended O-glycans (Fig. 5f). The jacalin used in the glycoproteomic approach to enrich for O-glycans does not allow the capture of these glycans (Fig. 5f). The reduction in ECM and cell surface derived peptides is therefore likely due to increased extension of O-glycans in KIC tumors compared to healthy mouse pancreas tissue.

### Glycoproteomic profiling of PDAC PDX samples reveals the conservation of GALA glycoprogram across mice and humans

Glycoproteomic analysis was performed on 18 PDX PDAC samples. Proteomic profiling identified approximately 4500-5500 proteins per sample, with a total of 6050 unique proteins detected across all samples (Fig. 6a). PCA at the protein level showed strong clustering of replicates (Fig. S6a). Glycopeptide analysis identified 1707 unique glycopeptides across the dataset, with more variability across samples compared to the proteome (Fig. 6b, Supplementary Table 8). Sample PDAC054T exhibited the highest glycopeptide count (∼1300), whereas PDAC021T and PDAC089T had the lowest (∼240 each). PDAC021T and PDAC089T contained ∼75 glycoproteins whereas other samples had 130-175 glycoproteins with 188 unique glycoproteins in total (Fig. 6c). PCA of the glycopeptide and glycoprotein datasets indicated high replicate consistency and that PDAC089T and PDAC021T were distinct from the other PDX samples, specifically at the glycopeptide level (Fig. 6d, S6b).

**Fig. 6.**
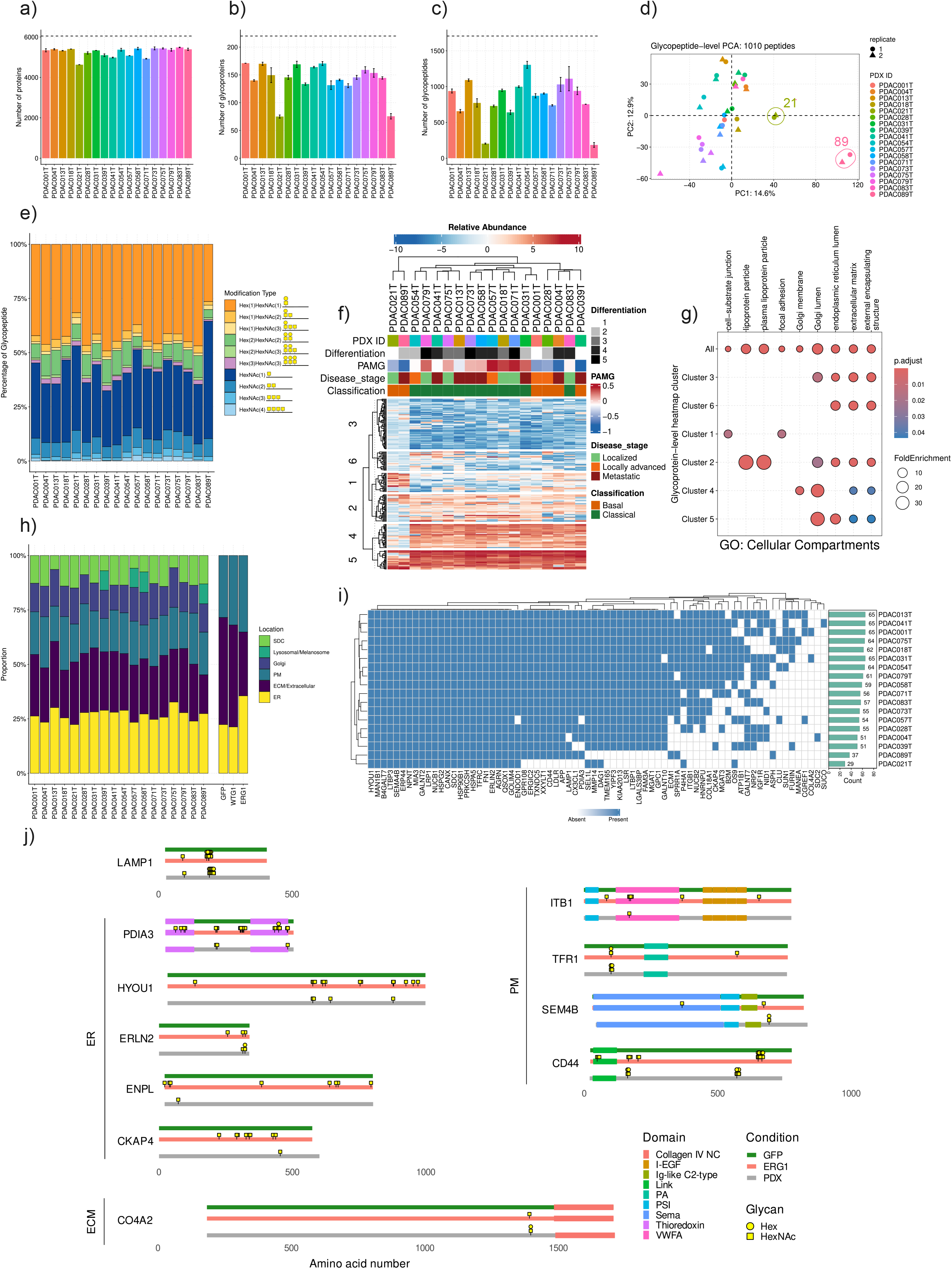
Glycoproteomic profiling of PDAC PDX tumours reveals inter-patient heterogeneity and overlap with ER-GALNT1-driven glycosylation signatures. **a,b,c.** Number of proteins, glycopeptides and glycoproteins detected in the analysis of 18 PDX PDAC samples. **d.** Glycopeptide-level PCA analysis. A total of 1707 unique glycopeptides were detected. **e.** Proportion of glycopeptide modifications detected per sample. Stacked bar plot showing each modification percentage per sample. **f.** Glycoprotein-level heatmap showing relative abundance comparison between PDX PDAC samples. K-means clustering was performed on protein rows and Euclidean distance on sample columns. Annotations shown include differentiation levels 1-5, pancreatic adenocarcinoma molecular gradient (PAMG) from -1 (orange) to 0.5 (green), basal (blue) or classical (yellow) classification, surgery status for resection (pink) or biopsy (blue) and disease stage gradient from localised to metastatic. **g.** Gene ontology cellular compartment analysis of each glycoprotein-level heatmap cluster. False discovery rate (p.adjust) is represented as a colour gradient from and dot size represents the fold enrichment. **h.** Comparison of top enriched gene ontology cellular compartment terms for the 18 PDX PDAC samples and comparison to cell line analysis of KPC47 GFP, WT-G1 and ER-G1. **i.** Heatmap representing presence (blue) or absence (white) of ER-G1 regulated glycoproteins in PDX samples. **j.** Linear glycosite maps of 11 ER-G1-regulated glycoproteins (log₂FC ≥ 2, adjusted p-value < 0.05) detected in PDX samples. Each protein displays three tracks (GFP, ER-G1, and PDX, top to bottom), positioned according to UniProt sequence annotations. Yellow squares denote detected glycopeptide sites. UniProt domains are color-coded and labeled.

In terms of glycans, all patients exhibited both T and Tn structures (Fig. 6e). Notably, PDAC089T, with the lowest number of glycopeptides detected, showed a disproportionately high proportion (56.87%) of HexNAc(1), suggesting a bias toward unextended Tn glycans. Hierarchical clustering analysis performed at the protein level revealed no clear clustering (Fig. S6c). By contrast, glycopeptide clustering revealed a separation of PDAC089T and PDAC021T from other PDXs (Fig. 6f). These PDX have previously been analysed at the transcriptomic level and classified using a newly defined metric, the Pancreatic Adenocarcinoma Molecular Gradient (PAMG), that aims to reflect tumour differentiation and aggressiveness^22^. However, this transcriptomic analysis did not segregate the two PDX, which appeared as basal types, with high PAMG. This suggests that glycopeptide signature could be used to better classify tumour types.

These two PDX had an enrichment for glycopeptides from proteins involved cell-substrate junction and focal adhesion, included proteins such as VIM (vimentin), PLEC (plectin) and PLAU (plasminogen activator, urokinase) (Fig. 6g, S6d,e). Cluster 5 was observed across all PDX samples and enriched for proteins associated with the ER and Golgi lumen. Three glycopeptide-level clusters showed enrichment for ER-related GO CC terms, indicating a strong ER signature (Fig. 6g, S6d,e). In fact, when the proportion of proteins with an ER-related GO CC was compared between PDX, they all appeared to contain a significant proportion, for most of them higher than in the control cell line KPC47 (GFP-KPC47) (Fig. 6h). Additionally, a large number of proteins belong to the extracellular matrix (ECM) and extracellular region, suggesting enhanced O-glycosylation affects the proteins secreted by PDAC cancer cells (Fig. 6h). Glycoproteins whose glycosylation was most strongly affected by ER-G1 expression in KPC47 (240 proteins with log₂ fold-change ≥ 2, adjusted p ≤ 0.05) were interrogated in PDX data. 55 ER-G1-regulated glycoproteins were detected across patient-derived xenograft (PDX) samples (Fig. 6i, Supplementary Table 9). Notably, glycoproteins shared across all PDX models included five ER-resident proteins: HSPA5, CANX, ERLIN2, MIA3, and HSP90B1.

To verify whether PDX and ER-G1 KPC47 shared glycosylation sites on specific proteins, glycopeptide positions were mapped onto UniProt protein sequences for GFP, PDX, and ER-G1 conditions with representative examples from ER-resident, cell surface, and ECM proteins shown (Fig. 6j). Remarkably, despite the significant evolutionary divergence, mouse and human proteins were glycosylated with very similar patterns and PDX-associated patterns resembled those obtained in ERG1 expressing cells (Figure 6j).

To explore whether the low glycopeptide levels of PDAC089T and PDAC021T could be attributed to expression of elongation enzymes as in KIC mouse tumors, the expression of O-glycosylation pathway enzymes was interrogated, but no pattern specific for 89 and 21 emerged (Fig. S6f). On the contrary, the expression of O-glycosylation enzymes appeared relatively conserved. Collectively, these findings indicate a widespread ER-associated O-glycosylation pattern across PDXs, with two samples displaying a unique glycopeptide pattern.

### The glycoprogram in PDAC tumors is multilayered

To explore the 220 glycoproteins detected across the PDX, their cellular compartment gene ontology was mapped and curated as for the KPC47 dataset. The map obtained contained ER, Golgi, plasma membrane, ECM and secreted proteins (Fig. S6g). While the specific proteins were sometimes different, similar biological processes appeared affected in both models. For instance, ECM adhesion proteins such as integrin Beta 1 (ITGB1), CD44, ECM remodelling factors MMP14, MMP1 and CANX were targeted (Fig. S6g). ECM proteins were highly represented. By contrast with KPC47 data, Insulin growth factor signaling is impacted by glycosylation, with IGF2, IGFR1 and four IGF binding proteins (IGFBP) represented (Fig. S6g). The IGF signaling axis has been implicated in PDAC progression and metastasis and individual proteins considered as therapeutic targets ^23^.

These data indicate that proteins in different subcellular locations and involved in cancer progression through different mechanisms are modified by O-glycosylation. Altogether, these results suggest GALA activates a program where multiple proteins are affected in coordinated fashion.

### ER-specific glycosylation targets structurally buried residues

Why does ER relocation of GALNT enzymes increase glycosylation of surface and secreted proteins, which normally transit through the Golgi and encounter Golgi-localised GALNTs? To address this question, glycopeptides were filtered to establish unambiguous glycosites, based on a single acceptor residue and/or available site-specific data ^24,25^. Corresponding protein 3D structures were obtained from AlphaFold2, and glycosites were analysed for solvent accessibility (Fig. 7a). A normalised, relative solvent-accessible surface area (rSASA) score was calculated for each amino acid in the identified proteins. Glycosites in KPC47 GFP cells generally scored above 0.6 (median 0.68), indicating high solvent exposure and suggesting that Golgi-localised GALNTs predominantly access the surface of proteic domains (Fig. 7b). By contrast, glycosites specific to KPC47 ER-G1 cells had significantly lower SASA scores (median 0.47), closely matching the SASA distribution of all serine and threonine (S/T) residues (Fig. 7b,c). This indicates that ER-localised GALNT1 glycosylates residues independently of structural constraints, consistent with co-translational glycosylation. Glycosites from healthy pancreas, KIC, and PDX samples were then analysed in the same way. Significantly lower SASA scores were observed in KIC tumours and human PDX samples than in healthy pancreas (Fig. 7d), with two PDX outlier samples (21 and 89) showing markedly lower median SASA scores than the rest.

**Fig. 7.**
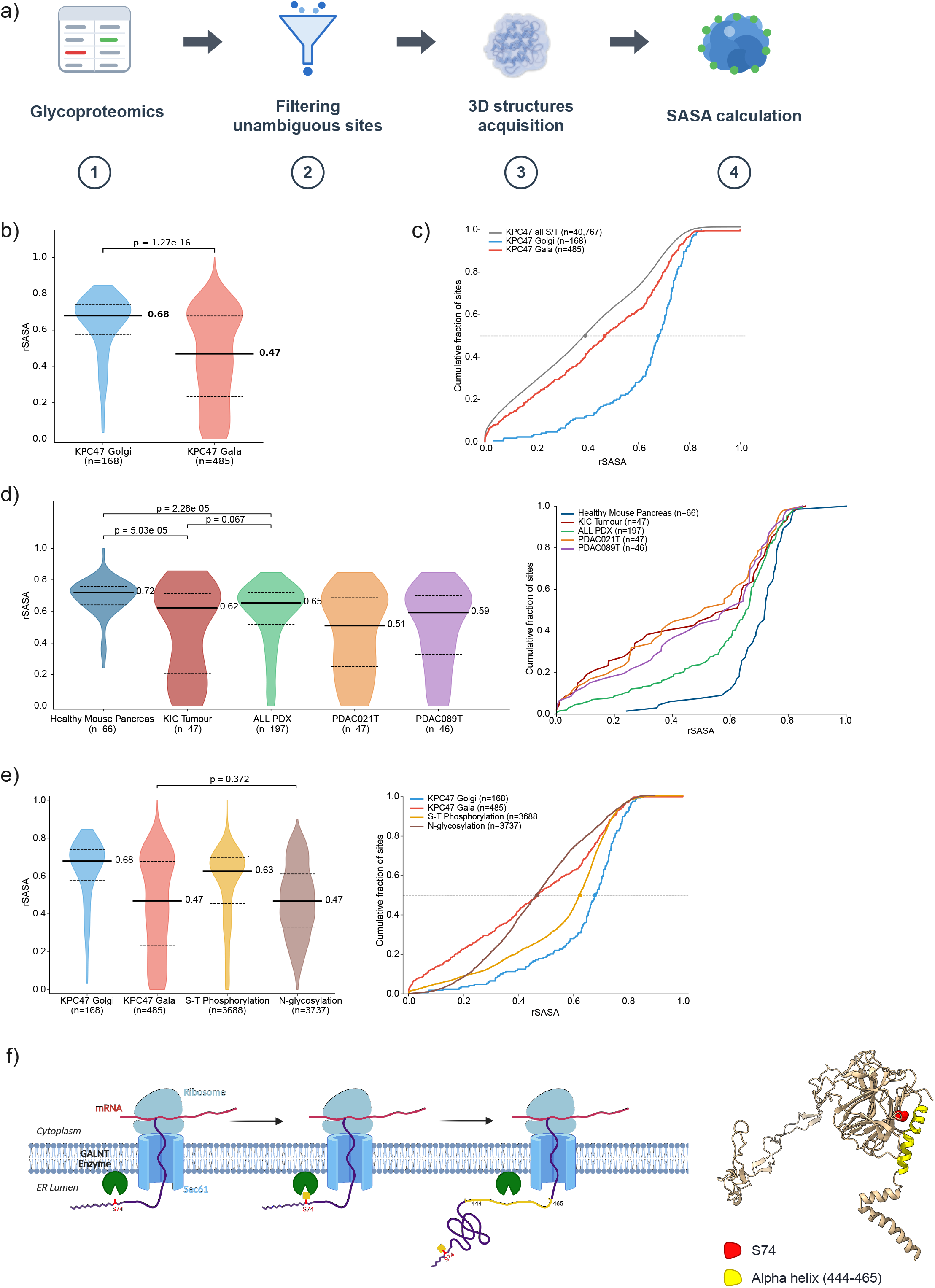
a. Schematic of SASA (Solvent-Accessible Surface Area) score generation. **b.** Violin plot of all unambiguous sites detected in KPC47 wild-type and in ER-G1 datasets. **c.** Cumulative fraction of sites in relation to rSASA score **d.** Violin plot and cumulative fraction of datasets obtained from tumors. **e.** Comparison of glycoproteomic data on the same set of proteins (n) with public data available for Ser/Thr phosphorylation and N-glycosylation. **f.** Calnexin synthesis and glycosylation, illustrating how a buried residue, Ser 74, could be glycosylated.

We then compared the SASA score distributions of two other well-known post-translational modifications — Ser/Thr phosphorylation and N-glycosylation — across all proteins in the dataset. Strikingly, the phosphorylation distribution suggested that kinases act mainly on exposed sites (median 0.67), whereas N-glycosylation, known to occur co-translationally, showed an rSASA median and distribution closely resembling that of GALNT-specific sites (median 0.47) (Fig. 7e).

To better understand the glycosylation of buried residues, calnexin, an ER-resident protein glycosylated and regulated by GALA was further analysed ^16^. The glycosylated Serine 74 (S74) is particularly buried within the main globular domain, with an rSASA of 0.08. This globular domain is formed by both N-terminal and C-terminal residues, with residues 444–465 forming an alpha helix that caps the globular domain (Fig. 7f). This configuration suggests that S74 is glycosylated before the alpha helix is formed and completes the globular domain. Overall, these results demonstrate that the relocation of GALNTs to the ER enables co-translational glycosylation on sites that remain cryptic when the enzymes are confined to the Golgi.

## Discussion

Our results demonstrate that spatial control of GALNTs is a major regulator of O-glycosylation in PDAC. In both PDAC biopsies and PDXs, Tn and GALNT2 frequently displayed ER-associated staining patterns, in marked contrast to healthy pancreatic tissue, where Tn and GALNTs are confined to the Golgi apparatus. The ER is a larger membrane system than the Golgi and distributed throughout the cell, explaining the change in intensity and appearance of Tn staining. Tn-modified calnexin, an ER-resident protein, is detected in numerous PDX samples but virtually absent from healthy pancreas. Finally, the substantial presence of glycosylated ER-resident proteins in the PDX and KIC glycoproteomes further supports activated relocation in both murine and human tumors. Our findings challenge the model that elevated Tn expression results from lack of elongation, due to loss of core-1 synthesis^8,26–28^. Transcriptomic and proteomic analyses of PDX samples revealed no consistent downregulation of C1GALT1 or COSMC, the Tn elongation factors, and glycoproteomic data identified the C1GALT1 product, the T glycan, in all PDXs analysed.

Beyond initiation, O-glycans structures are further shaped by elongation and branching enzymes that are themselves altered in cancer ^6,29^. In our study, KIC tumors transcriptionally upregulate core-2 synthesis enzymes GCNTs. Upregulation of GCNT1 has been implicated in prostate cancer, suggesting GCNTs could also promote pancreatic tumor growth in some cases ^30^. The elongation of O-glycans could also explain why some of the PDX have a low Tn glycopeptide abundance and represent a challenge for future glycoproteomic efforts.

PDAC tumors display inter- and intra-tumoural heterogeneity. Tn was prominently detected in CK19⁺ ductal tumour cells, but CK19⁻/Tn⁺ cells were also present across multiple samples. Although the identity of these cells remains uncertain, this suggests that ER O-glycosylation is not confined to the epithelial compartment. The frequency of spatial reorganisation of O-glycosylation in PDAC suggests that, as in hepatocellular carcinoma, it is a driver of tumor growth ^15^. Consistently, ER-glycosylation hyperactivation in an orthotopic model led to marked stimulation of tumor growth, while GALA inhibition blocked its development.

The quantitative glycoproteomics analysis reveals how O-glycosylation reorganisation generates a different glycoproteome in affected cells. Many glycosites detected under ER relocation were undetectable in the reference cell line, contrasting with the limited effect of GALNT1 overexpression. In fact, endogenous GALNT1 expression appears sufficient to saturate glycosylation sites in the Golgi.

How to then explain the dramatic increase linked to the ER? GALNTs possess lectin domains that bind GalNAc residues and promote neighbouring site glycosylation. In the Golgi, elongation of Tn by C1GALT and other enzymes can compete with this lectin-driven glycosylation. In the ER, the absence of elongation enzymes allows accumulation of poly-Tn structures. PDX samples displayed abundant poly-Tn structures and extensive modification of ER-resident proteins. In the two PDXs with lower glycopeptide abundance, ER-resident proteins remained well represented, suggesting that reduced glycopeptide levels reflect tumour heterogeneity or differences in glycan maturation rather than absence of GALA activity. Similar to KIC mouse tumors, some human tumours may generate more extended O-glycan structures, which could mask Tn and T antigens in the lectin-based enrichment strategies. Such shifts toward more elaborated glycoforms have been associated with aggressive tumour behaviour and may represent adaptive mechanisms for immune evasion or altered cell–ECM interactions ^31,32^. For instance, the core-2 forming enzyme GCNT1 has recently been shown to promote prostate tumor growth ^33^. Thus, in tumors, the stimulation of O-glycosylation initiation by GALA is probably combined with other processes in the O-glycan synthesis pathway.

A central implication of our data is that GALA converts O-glycosylation from a post-translational into a co-translational modification. In healthy cells, O-GalNAc glycosylation is initiated in the Golgi, after proteins have folded and assembled — in contrast to N-glycosylation, which is added co-translationally in the ER ^34,35^. Newly synthesised proteins emerge from the Sec61 translocon into the ER in an unfolded, extended state, where they are modified by the oligosaccharyltransferase (OST) complex. By relocating GALNTs to this compartment, GALA likewise renders O-glycosylation co-translational, allowing the enzymes to reach residues that become inaccessible once folding and complex assembly are complete. For example, Calnexin is glycosylated on a site (S74) that is deeply buried in the fully folded protein.

The low SASA of N-glycosylation sites arises because these values are computed from structures of unglycosylated proteins, typically expressed in bacteria: the protein folds without the glycan, burying a site that is modified co-translationally in mammalian cells. The same logic applies to the buried O-glycosylation sites we identify. A corollary is that addition of a glycan at such a site will affect the structure of the protein. It is well documented that N-glycans influence protein conformation — as demonstrated, for instance, in the Fc domain of IgG ^36–38^. We expect that O-glycosylation of buried residues similarly induces conformational changes relative to the unglycosylated form. The relocation of GALNTs could thus represent a powerful mechanism to regulate and expand cell surface activities by generating alternative glycoforms with modified conformations.

Previous work has shown that GALA promotes ECM degradation through glycosylation of two proteins, MMP14 and CANX, which act respectively as a protease and a disulfide bond reductase ^15,16^. GALA glycosylation thus coordinates enzymatic activities. Beyond ECM degradation, numerous proteins implicated in PDAC progression were hyperglycosylated ^39^. Altered glycosylation of ER-resident proteins may additionally influence protein folding and ER homeostasis, processes increasingly recognised as critical for tumour cell survival ^40^.

In cancer, transcriptional programs such as the epithelial-mesenchymal transition or the MYC-driven proliferation program have been recognized and studied ^41,42^. Similarly, kinase-mediated programs, such as the DNA damage response coordinated by the ATM kinase, allow cells to coordinate activity of multiple proteins to respond to signals or challenges ^43^. Our results suggest that spatial control of O-glycosylation initiation functions as a regulatory switch, allowing GALNTs to access cryptic sites in numerous proteins and initiate a “glyco-program” that coordinately modulates the activity of multiple enzymes, receptors and adhesion molecules. The glycoproteomic dataset presented here provides a foundation for exploring how this O-glycosylation program contributes to tumour progression and offers new opportunities to identify biomarkers and therapeutic targets in PDAC.

## Methods

### Patient samples, cell lines and mouse tumor samples

Human tissue microarrays (TMA) containing human PDAC and normal tissue were obtained from Biomax (Catalogue number PA482). Patient-derived xenograft (PDX) tumours were derived from the PaCaOmics study (2011-A01439-32). The murine KPC47 cell line is a mouse pancreatic ductal adenocarcinoma (PDAC) cell line derived from the KPC genetically engineered mouse model (LSL-Kras^G12D/+; LSL-Trp53^R172H/+; Pdx1-Cre) and supplied by David Tuveson (Cold Spring Harbour Laboratories, NY, USA). Cells were maintained in Dulbecco’s Modified Eagles Medium (DMEM; D5796; with 4500 mg/L glucose, L-glutamine, and sodium bicarbonate, without sodium pyruvate; Sigma-Aldrich) supplemented with 10% (v/v) foetal bovine serum (Life Technologies) at 37°C and 5% CO_2_. The KPC47 cell line was engineered to stably express an empty vector, ER-localised GALNT1, Wild-type GALNT1, ER-2Lec and a catalytic domain mutant ER-GALNT1 using the Sleeping Beauty transposon system.

### PDAC mouse models

For PDAC tissue, male Pdx1-Cre; Ink4a/Arf fl/fl; LSL-KrasG12D mice, developing spontaneous PDAC, between 8 and 12 wk of age, and their mating control littermates were used, as previously described ^44^. For orthotopic graft of KPC47 cell lines, 10,000 KPC47 cells in 20 µl of ice-cold matrigel were implanted into the pancreas of C57/Bl6 mice, six to eight weeks old. Mice were administered doxycycline in drinking water at 2mg/ml with 2% sucrose and via intraperitoneal injection twice a week at 50mg/kg for the duration. Mice were euthanised at six weeks post-injection and organs harvested for histology. Orthotopic protocols were reviewed and approved by the Institutional Animal Care and Use Committee (IACUC) of the Agency for Science and Technology and Research in Singapore.

### Antibodies and reagents

Biotinylated Vicia villosa lectin (VVL, B-1235, Vector Laboratories), Alexa Fluor 647 Helix pomatia lectin (HPL, Invitrogen, L32454), anti-cytokeratin-19 (CK19, Proteintech, 10712-1-AP), anti-GRP78 (Proteintech, 66574-1-Ig), anti-GALNT2 (Sigma, HPA011222), anti-GOLPH2 (Proteintech, 66331-1-Ig), Hoechst (Thermo Scientific, 62249), anti-calnexin (abcam, ab22595), anti-GAPDH (Proteintech, 60004-1-Ig), anti-α-tubulin (Invitrogen, 62204), anti-V5 tag (Invitrogen, R960-25), anti-GALNT1 (Abcam, ab253025), VVL-agarose (Vector Laboratories, AL-1233-2), Jacalin-agarose (Thermo Scientific, 20395), anti-CD29 (9EG7; BD Biosciences, 550531), anti-paxillin (BD Biosciences, 610051), anti-FAK (Proteintech, 12636-1-AP), Phalloidin-488 (Biolegend, 424201). Matrigel solution (354230, Corning).

### Immunofluorescence

Tissue samples were deparaffinised in xylene and rehydrated by decreasing concentration of ethanol to water washes. Heat-induced epitope retrieval was performed using pH6 citrate buffer (RE7113-49 CE; Leica Biosystems). Tissue sections were blocked for 1 hour with 1% (v/v) Triton X-100 and 6% (v/v) goat serum in 1x PBS. For samples that were incubated with lectin-biotin reagents, an initial streptavidin-biotin block was used to block endogenous biotin (SP-2002; Vector Laboratories). Samples were then incubated overnight at 4°C with primary antibodies and lectins diluted in 1% (v/v) Triton X-100 and 1% (v/v) goat serum in 1x PBS. Three wash steps were performed in 1x PBS before incubation for 1 hour at room temperature (RT) with secondary antibodies or streptavidin-conjugated fluorophores diluted in 1% (v/v) Triton X-100 and 1% (v/v) goat serum in 1x PBS. Then samples were washed three times in 1x PBS and nuclei were stained with Hoechst 33342 (1:1000; ThermoFisher) in dH20 for 5 minutes at RT. Samples were then washed in 1x PBS followed by dH20, before mounting coverslips (Antifade Mounting Medium; H-1000-10; Vectashield) and sealing with nail polish. Mounted slides were stored at 4°C and protected from light before confocal imaging. Images were captured using an ImageXpress® Confocal HT.ai High-Content Imaging System with a 20x water objective. For immunofluorescence of cells, the HPL area per cell was analysed using MetaXpress analysis software.

Analysis of tissue microarray cores was performed using QuPath (0.5.1). For VVL area the tumour core area was thresholded by selecting the DAPI area. The VVL area was defined by selecting all the strong stained VVL at a constant threshold above background levels. The VVL area was then quantified as a percentage of total core area (Fig. S1b). For the classification of CK19 and VVL-positive cells, all cells were selected and segmented using DAPI and expanding the cell region (Fig. S1c). Thresholds for CK19 and VVL were created to select the positive signal. Classifications were performed in order to allocate cells as VVL+CK19-, VVL+CK19+, VVL-CK19+ and VVL-CK19-. The values are given as a proportion of total cells within the tumour core.

### Immunohistochemistry

Samples were deparaffinised in Bond Dewax solution (AR9222; Leica Biosystems) and rehydrated with 100% (v/v) ethanol through to 1x Bond Wash solution (AR9590; Leica Biosystems). Samples were boiled for 40 minutes at 100°C using Bond Epitope Retrieval solution, pH6 (RE7113-CE; Leica Biosystems). Endogenous peroxidases were blocked with 3% (v/v) H2O2 for 15 minutes and incubated with 10% (v/v) goat serum block for 30 minutes. Samples were then incubated with a streptavidin-biotin blocking kit to block endogenous biotin (SP-2001; Vector Laboratories). VVL-biotin was added for 1 h at room temperature. After washing three times with Bond wash solution, samples were incubated with streptavidin HRP at room temperature for 30 minutes. After washing with Bond wash solution, Bond Polymer Refine Detection (DS9800; Leica Biosystems) mixed DAB was used to visualise HRP signals. Nuclei were then counterstained with hematoxylin for 5 minutes. Samples were dehydrated and mounted before slide scanning. Analysis was performed using QuPath (0.5.1). Cells were segmented using hematoxylin and VVL positivity selected by thresholding DAB (Fig. S1f). The area and number of VVL cells were quantified.

### RNA-sequencing and analysis

RNA-seq reads were mapped using STAR^45^ considering the ENCODE annotations. The content from human and mice was separated using the SMAP algorithm having the human hg19 and mouse mm38 genomes (Ensemble 75). Then, the gene expression profiles were obtained using the FeatureCount^46^ program. The heatmap was generated to observe the expression profiles for GALNTs, C1GALT1C1 and C1GALT1, using the gene counts obtained after being normalised using the upper-quartile approach^47^.

### VVL immunoprecipitation

PDX PDAC tumour samples, healthy mouse pancreas and cell-derived xenograft tumours from MDA-MB-231 (positive control) were homogenised in ice-cold RIPA lysis buffer and lysed for 1 h at 4°C with rotation. Samples were then centrifuged at 13,000 x g for 15 min at 4°C and the supernatant was collected. Lysates were incubated with VVL-agarose (Vector Laboratories) overnight at 4°C. The beads were washed with RIPA buffer three times and then eluted in 2x LDS sample buffer with 50 mM DTT. Samples were then boiled at 95°C for 10 min before SDS-PAGE electrophoresis and immunoblotting.

### Gel electrophoresis and immunoblotting

Lysates and VVL-immunoprecipitation (VVL-IP) elutions were separated by SDS-PAGE using 4-12% Bis-Tris NuPage gels (Invitrogen) at 170 v for 30 min. Separated proteins were transferred onto a 0.45 µm nitrocellulose membrane (GE Healthcare) using a Trans-Blot® Turbo™ Transfer System (Bio-Rad) according to the manufacturer’s instructions. Membranes were blocked with 5% (w/v) bovine serum albumin (BSA) in 1 x PBS for 1 h. For biotinylated lectins, Streptavidin/Biotin Blocking kit (SP-2002) was used to block non-specific binding as per the manufacturer’s instructions. Membranes were incubated with primary antibodies or lectins in 5% (w/v) BSA in tris-buffered saline with Tween 20 (1x TBST [0.05% (v/v) Tween 20, 1 mM Tris, 15 mM NaCl]) for 1 hour at room temperature or 4°C overnight, followed by secondary HRP-conjugated antibodies for 30 min. Membranes were visualised using chemiluminescence and images with an Amersham ImageQuant 800 system (Cytiva).

### Glycoproteomics and proteomics sample preparation

Cell pellets were homogenised in 400 µl of 50 mM ammonium bicarbonate (AmBic) and placed in a water bath sonicator for 30 min with ice. Approximately 5-10 mg of frozen PDX tumour tissue was homogenised on a bead mill in 250 µl 50 mM AmBic with stainless steel beads. The PDX tissue was then placed in a Covaris sonnicator (Sonolab) using microTUBE-500 tubes (Covaris, 520185) at 80-100W PIP, 10% duty factor and 200 cycles per burst for 180 s with continuous degassing. Protein concentration was assessed by the Pierce BCA Protein Assay. For DDA libraries 500 µg of lysate and DIA 200 µg was further processed.

Volumes were adjusted to 400 µl, followed by reduction at 60°C for 45 min in 5 mM dithiothreitol (DTT) and alkylation in darkness for 30 min at room temperature. Alkylation was terminated by 10 mM iodoacetamide with 5 mM DTT for 15 min. Then, 0.1 U neuraminidase from *Clostridium perfringens*, (Sigma, N2876) was diluted in sodium acetate buffer (0.1 M, pH 5) and incubated at 37°C for 2 h. The interaction was stopped by 10 min incubation at 98°C, before cooling on ice. Samples were then digested in 1:50 Sequencing Grade Modified Trypsin (V511A, Promega) for 60 min at 47°C and 10 h at 37°C. Digests were acidified with trifluoroacetic acid (TFA), centrifuged 21,000 xg for 30 mins and supernatant retained. Peptides were lyophilised in a SpeedVac.

### Glycopeptide sample enrichment

Glycopeptide samples for the DIA and DDA analysis were enriched according to sLWAC-HTP protocol. Briefly, sLWAC-HTP protocol is the further development of sLWAC protocol from Glyco-DIA method^17^ towards its miniaturisation and high-throughput capability (to be published elsewhere). Briefly, previously desialylated peptide samples were resuspended in sLWAC loading buffer (175mM Tris Buffer, pH-7.4), incubated with Jacalin lectin (conjugated on agarose) beads, washed (175mM Tris Buffer, pH-7.4) and eluted with 175mM Tris Buffer containing 0.8M galactose, pH-7.4.

### LC–MS/MS method

EASY-nLC 1200 UHPLC (Thermo Fisher Scientific) interfaced via nanoFlex ion source (Thermo Fisher Scientific) to an Orbitrap Fusion/ Lumos Mass Spectrometer (Thermo Fisher Scientific) was used for all the MS analyses. The nano-flow liquid chromatography (nLC) was operated in a single analytical column set up with PicoFrit Emitters (New Objectives, 75 μm inner diameter) packed with Reprosil-Pure-AQ C18 phase (Dr. Maisch, 3 μm particle size, 19–21 cm column length).

### LC–MS/MS DDA method

Each sample was injected onto the column and eluted in a 2 h gradient from 3% to 32% B in 95 min, from 32% to 100% B in 10 min and 100% B for 15 min, at 200 nl min−1 (solvent A, 100% H2O; solvent B, 80% acetonitrile; both containing 0.1% (v/v) formic acid). A precursor MS scan (m/z 350–1,700) of intact peptides was acquired in the Orbitrap at the nominal resolution setting of 120,000, followed by Orbitrap HCD–MS/MS at the nominal resolution setting of 50,000 of the 15 most abundant multiply charged precursors in the MS spectrum; a minimum MS signal threshold of 20,000 was used for triggering data-dependent fragmentation events. HCD collision energy of 37% ± 5% was used in all the DDA runs in the study. A 40 s dynamic exclusion window was used to prevent repeated analysis of the same components.

### DDA data analysis and library generation

The DDA data processing was performed using Proteome Discoverer (PD 2.4) software using Sequest HT as previously described^17^ with some changes. Briefly, all spectra were searched with full and semi-specific enzymatic cleavage. In all cases, the precursor mass tolerance was set to 15 ppm and fragment ion mass tolerance to 20 milli mass unit (mmu). Carbamidomethylation on cysteine residues was used as a fixed modification. Methionine oxidation, HexNAc and Hex-HexNAc attachment to serine, threonine and tyrosine were used as variable modifications for MS/MS. All spectra were validated by Percolator Node with the false discovery rate of 1% and additionally filtered by XCore >1.2. Spectral libraries were generated as previously described^17^. Briefly, the acquired DDA result files (Thermo’s Mass Spec Format, pdResult) were imported to Spectronaut using the spectral library generation functionality of Spectronaut with default settings. A peptide was added to the spectral library, containing at least 3 and at most 8 of the most intensive peptide fragment ions. Supplementary table 10 contains an overview of the generated spectral libraries.

### LC–MS/MS DIA method

The mass spectrometry data has been deposited to the ProteomeXchange Consortium via the PRIDE repository with the dataset identifier PXD081077.

Each sample was injected onto the column and eluted in a 1 h gradient from 3% to 32% B in 40 min, from 32% to 100% B in 5 min and 100% B for 15 min, at 200 nl min−1 (solvent A, 100% H2O; solvent B, 80% acetonitrile; both containing 0.1% (v/v) formic acid).

The DIA method consisted of an MS scan from m/z 400 to 1,200 at the nominal resolution setting of 120,000 and targeted MS/MS scans with 40 windows of 20m/z isolation width windows to generate DIA segments from m/z 400 to 1,200. DIA segments were acquired at the nominal resolution setting of 30,000 with custom automatic gain control (AGC) target (1000% normalized). HCD collision energy of 37% ± 5% was used in all of the DDA runs in the study. DIA data were analysed with Spectronaut 18.0 (Biognosys AG) as previously described^17^. Briefly, all DIA glycoproteomics data were analysed using Glyco-DIA library based approach^17^ for KPC47 samples and library free direct DIA analysis for PDX samples. All DIA proteomics data were analysed using library free DIA approach using the default Spectronaut settings. For direct DIA analysis carbamidomethylation on cysteine residues was used as a fixed modification and methionine oxidation was used as variable modifications. For glycoproteomic direct DIA analysis, HexNAc and Hex-HexNAc attachment to serine, and threonine were additionally chosen as variable modifications. Decoy generation was set to inverse with the decoy limit of 5000.

### Bioinformatics and statistics

All bioinformatics and statistical analyses were performed using GraphPad Prism (10.4.0), R Statistical software (4.6.0) and Python (3.14.6). Quantitative proteomic and glycoproteomic data were processed in R using the Bioconductor packages limma (v3.68.2), vsn (v3.80.0) and SummarizedExperiment (1.42.0), and functional enrichment was performed with clusterProfiler (v4.20.0). For glycopeptide-level analysis, entries lacking essential identifiers, non-glycopeptides, and exact duplicates were removed. After variance-stabilising normalisation, missing values for ER-G1-specific glycopeptides in WT1/GFP samples were imputed by minimum probability imputation (Fig. S4c). Glycopeptides carrying more HexNAc residues than available Ser/Thr sites were discarded as biologically impossible (O-GalNAc occurs only on Ser/Thr). Because the glycan positions reflect a default placement rather than an experimentally scored localization, glycosites were considered unambiguous only when the number of HexNAc residues equalled the number of Ser/Thr sites, the single case in which placement is unique; ambiguous glycopeptides were retained and flagged for downstream site assignment. For the comparative analysis, sites significantly enriched in ER-G1 relative to control were defined as GALA sites; sites present in the control condition served as the Golgi reference. Filtered glycopeptide, glycoprotein and total-proteome intensities were assembled into SummarizedExperiment objects. Features were required to be quantified with no more than one missing value in at least one experimental condition. Intensities were log2-transformed and normalized by variance-stabilizing normalization (vsn). For the human PDX dataset, glycopeptide and glycoprotein intensities were additionally normalized to the matched total-proteome abundance to account for differences in underlying protein expression. Remaining missing values, which predominantly reflected low-abundance (missing-not-at-random) signals, were imputed using a left-censored MinProb model (q = 0.01) with a fixed random seed to ensure reproducibility. Differential abundance between conditions was assessed with linear models and empirical-Bayes moderated t-statistics (limma), using the contrasts ER-G1 vs GFP, ER-G1 vs WT-G1 and WT-G1 vs GFP for the KPC47 model and tumour vs healthy pancreas for the KIC model. P-values were corrected for multiple testing across all features using the Benjamini-Hochberg procedure to control the false discovery rate (FDR). Unless otherwise stated, glycopeptides and glycoproteins were considered significantly regulated at an adjusted p-value <= 0.05 and an absolute log2 fold-change >= 2; for selected volcano-plot and heatmap analyses a more stringent threshold (adjusted p <= 0.01, |log2FC| >= 3) was applied. Over-representation analysis of Gene Ontology terms (cellular component, molecular function and biological process), KEGG pathways and Reactome pathways was performed with clusterProfiler, using org.Mm.eg.db (mouse: KPC47, KIC) or org.Hs.eg.db (human: PDX) as annotation databases. Gene identifiers were mapped to Entrez IDs, and the full set of proteins identified and quantified in the corresponding dataset was used as the statistical background (universe). Terms were considered enriched at a p-value cutoff of 0.05 after Benjamini-Hochberg correction. Enrichment across sample groups or co-expression clusters was summarized using compareCluster. To compare the mouse KPC47 and human PDX glycoproteomes, mouse gene symbols were mapped to their human orthologs with the homologene package (mouse, taxid 10090 -> human, taxid 9606); genes without an assigned ortholog were matched by their upper-cased symbol. Graphics were generated using ggplot2(v4.0.3)[40] and ComplexHeatmap (v2.28.0). For heatmaps, row-wise features were segmented by K-means clustering, and samples were grouped by hierarchical clustering with Euclidean distance (Ward.D2 linkage for the PDX heatmaps; complete linkage for the mouse heatmaps). UniProt protein-feature annotations were queried with drawProteins.

### Proteomic data candidate selection and network analysis

Glycosylated proteins were identified from KPC (Mus musculus) and PDX (Homo sapiens) analyses. Differentially glycosylated candidates in KPC data were selected as proteins up-regulated in ER-G1 relative to GFP (log2FC >= 2 and adjusted p-value <= 0.05; 229 proteins). PDX candidates comprised all identified glycosylated proteins (220 proteins). Proteins were classified into subcellular compartments (ER, Golgi, plasma membrane, extracellular matrix, nucleus, lysosome, cytoskeleton) using automated rules based on GO Cellular Component annotations and transmembrane domain annotations retrieved from UniProt (taxonomy IDs 10090 and 9606, accessed September 1, 2025). Final compartment assignments were manually reviewed and adjusted (Fig. S4k). PPI networks were constructed using the STRING database (v12.0; confidence score >= 0.4), queried with UniProt accession IDs via the Cytoscape v3.10.4 STRING app, independently for Mus musculus and Homo sapiens. Full compartment networks were arranged using the Attribute Circle layout based on subcellular compartment assignments, with nodes colored by compartment category, manual repositioning, and edge bundling applied using default parameters. For the KPC plasma membrane subnetwork (n = 65), functional enrichment data were overlaid using the STRING Enrichment underlay and the top 5 enriched GO Biological Process and KEGG terms were visualized using the yFiles Tree layout.

### Structural accessibility analysis

For each protein with an identified glycosite, the corresponding AlphaFold2 structure was retrieved via its UniProt accession. Per-residue solvent-accessible surface area (SASA) was calculated with FreeSASA (default parameters) and normalised to relative SASA (rSASA, 0–1) using residue-type theoretical maxima from Tien et al. (2013). rSASA values were extracted for each glycosite, with all S/T residues of the same proteins used as background reference. Ser/Thr phosphorylation and N-glycosylation sites annotated on these proteins (UniProt) were processed identically for comparison. rSASA distributions were compared using two-sided Mann-Whitney U tests, with cumulative distribution functions used to visualise the score ranges. All analyses were performed in Python 3.12.

### Structural visualisation

The AlphaFold2 model of Calnexin was examined in ChimeraX to localise glycosylated Ser74 and the hydrophobic α-helix (residues 444–465) within the globular domain.

## Supporting information

Tables of glycosylated proteins

## Author contributions

R.B. designed and performed experiments and ran analyses

E.L. assisted bioinformatics analyses, assembled figures and wrote text

G.D.R. ran bioinformatic analysis

X.L.G. performed experiments and provided intellectual input

M.K. performed experiments

E.M. provided samples and feedback on the manuscript

N.D. provided human samples and feedback on the manuscript

A.C. provided transcriptomic analyses

R.T. provided murine samples and feedback on the manuscript

S.V. provided murine samples and feedback on the manuscript

F.G. managed murine sample collection and storage

B.C. provided human samples and feedback on the manuscript

M.J.H. provided support and feedback on the manuscript

S.V. performed mass spectrometry experiments and analyses

F.B. designed experiments, coordinated the study and wrote the manuscript

## Acknowledgements

F.B. is supported by the CNRS and grants from AMIDEX (AMX-20-CE-03) and by Leader in Oncology grant from the “Fondation ARC pour la recherche sur le cancer”. S.Y.V is supported by the Danish National Research Foundation (DNRF196). Thank you to Marion Rubis for providing the PDX immunohistochemistry data.

## Competing interests

F.B. is a co-founder, shareholder and CEO of Albatroz therapeutics Pte Ltd. X.L.G. is a shareholder of Albatroz therapeutics. Other authors declare no competing interests.

---

**Supplemental Figure 1.**
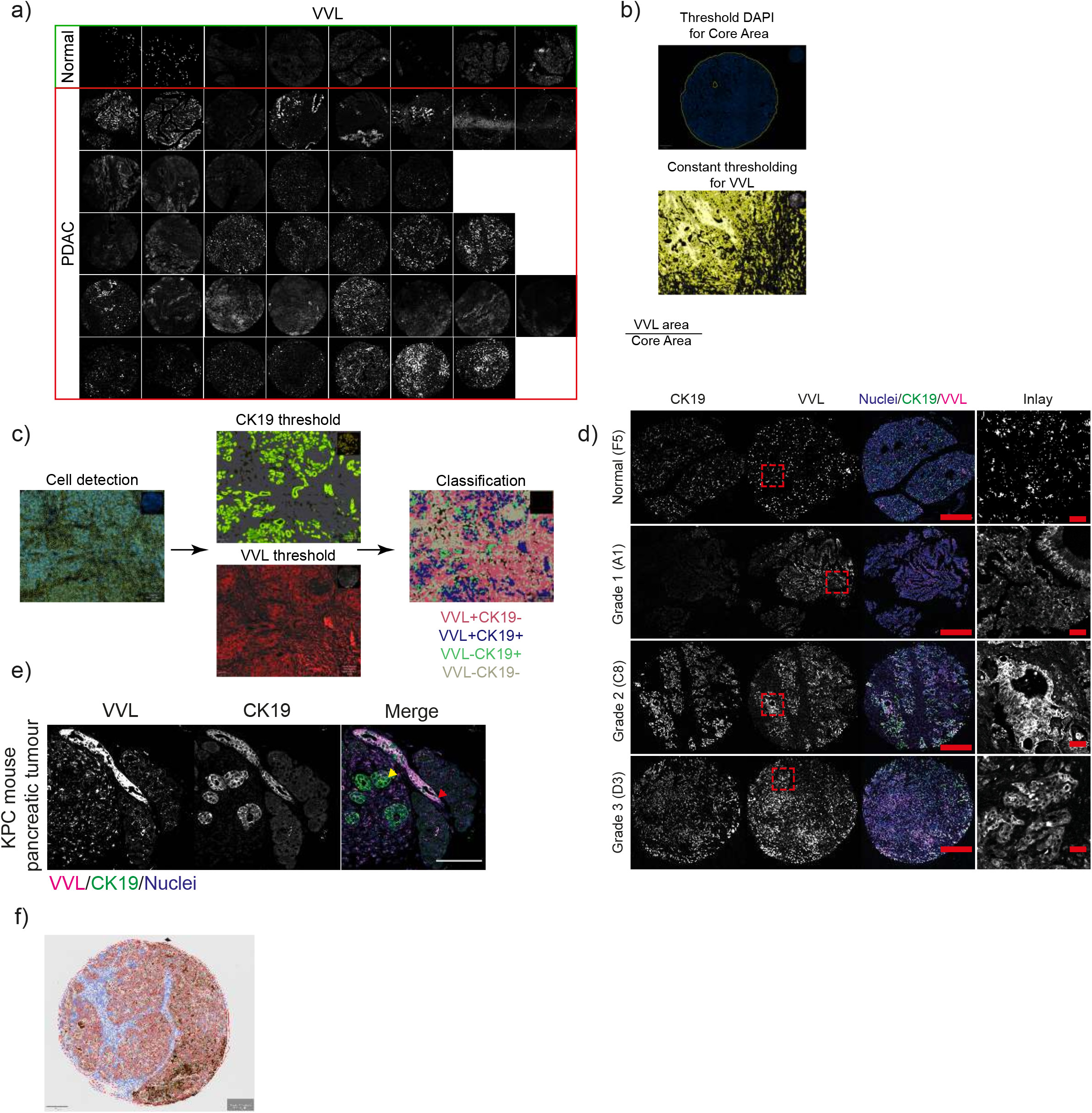
**a.** Immunofluorescence staining of Tn glycan using VVL on human TMA sections from pancreatic ductal adenocarcinoma (PDAC) and normal tissue. **b.** VVL area quantification method for TMA analysis. **c.** Analysis pipeline performed for CK19 and VVL classifications. Cells were detected by nuclei and cell expansion. Thresholds for CK19 and VVL staining were performed and based on the cell mask and threshold, each cell was given a classification based on CK19 and VVL positivity. **d.** Representative images showing CK19 (green), Nuclei (blue), and VVL (magenta) for normal tissue and multiple PDAC cores within the TMA from different grades. Individual channels are shown in white (scale bar: 300 µm). The inlay displays a zoomed region of VVL staining (scale bar: 50 µm). **e.** Immunofluorescence of KPC PDAC tumour tissue for VVL (magenta) and CK19 (green). CK19+VVL+ region is shown by red arrow and CK19+VVL- is shown by yellow arrow (scale bar 50 µm). **f.** Cell masking for PDX VVL immunohistochemistry analysis shows the classification of VVL positive (red) and negative cells (blue).

**Supplemental Figure 2.**
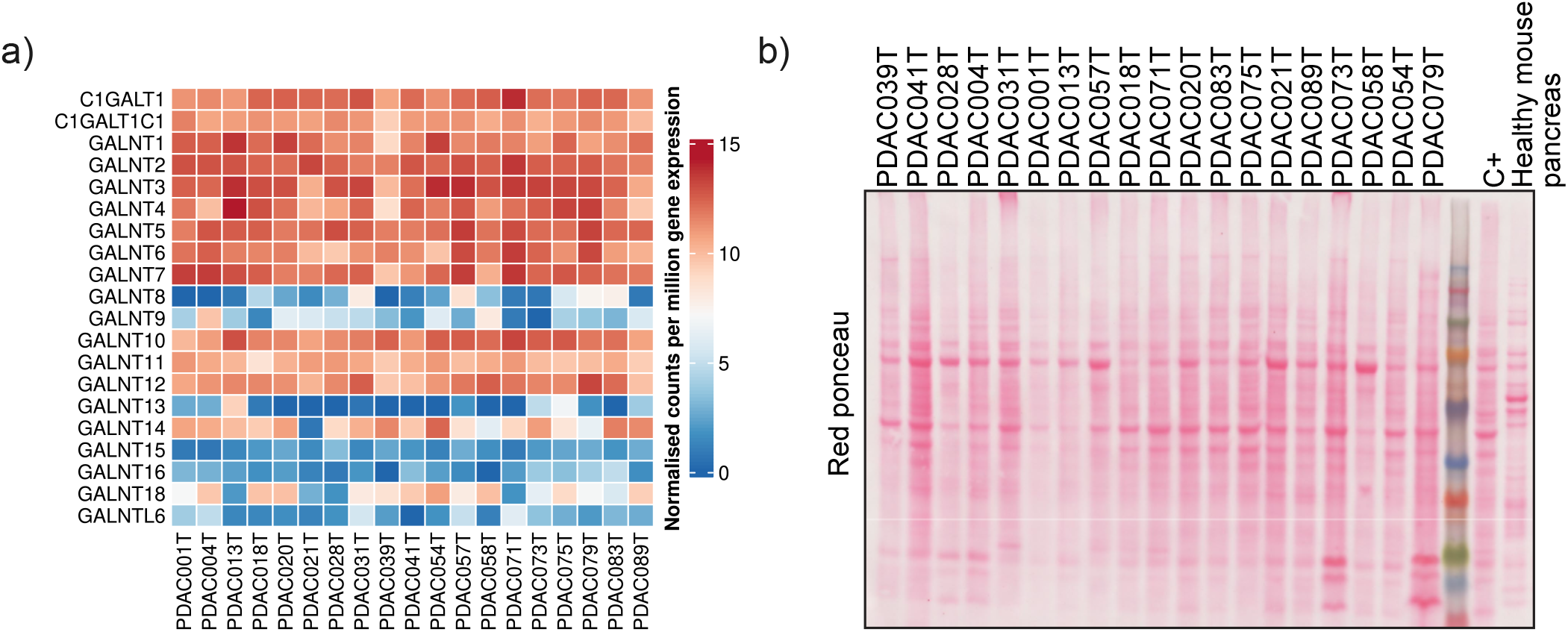
**a.** RNA-seq counts per million (CPM) gene expression of 19 PDX PDAC tumours for N-acetylgalactosaminyltransferases (GALNTs), C1GALT1 and C1GALT1C1. Expression levels are shown from 0 (blue) to 15 (red). **b.** Red Ponceau staining of the input samples used for VVL IP analysis described in Figure 2e.

**Supplemental Figure 3.**
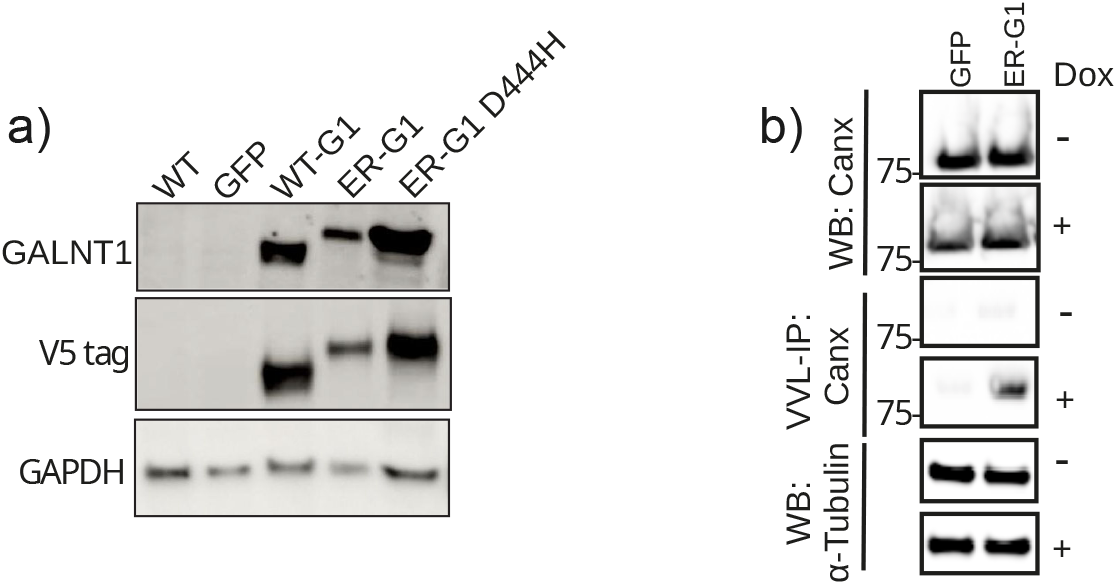
**a.** Immunoblot of KPC47 cell lysates for GALNT1, V5 tag and GAPDH. Wild-type (WT), expression of GFP, WT-G1, ER-G1 or ER-G1 H211D. **b.** VVL-agarose enrichment (VVL-IP) of GFP and ER-G1 lysates (WB) with immunoblot for Canx and ⍺-Tubulin.

**Supplementary Figure 4.**
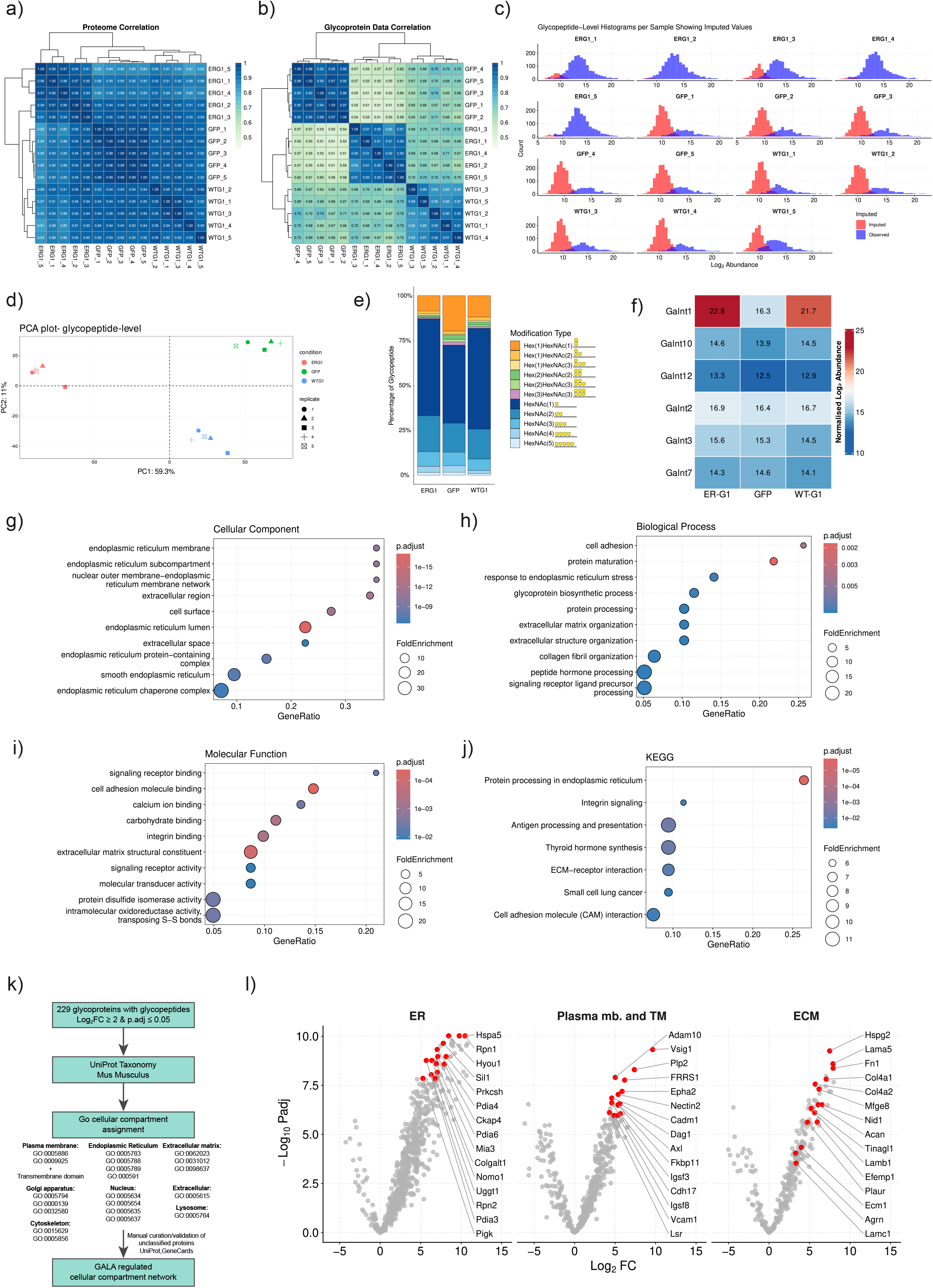
**a.** Coefficient of variation analysis of proteome abundances before variation-stabilising normalisation. Scale represented from low (light green) to high (dark blue). **b.** Coefficient of variation analysis of glycopeptide abundances before normalisation. **c.** Histograms per sample showing log_2_ abundance and peptide count. Observed data (blue) and minimum probability imputed values (red) are shown. **d.** Glycopeptide-level PCA plot of PC1 and PC2. **e.** Glycopeptide modification type per cell line as a percentage of total glycopeptides. **f.** Proteome-level abundance analysis of the GALNT proteins detected in KPC47. The average normalised Log_2_ abundance is shown for n=5. **g.** Schematic of subcellular compartment classification rules applied to candidate proteins. Compartment assignments were based on GO Cellular Component annotations and transmembrane domain annotations from UniProt, with manual expert review. **h,i,j,k.** Over-representation analysis was performed on the 240 proteins showing >4-fold increased (adjusted P-value <0.05) O-glycosylation in KPC47-ERG1 cells, assessing Gene Ontology **h.** Cellular Component **i** KEGG pathways, **j.** Molecular Function, **k.** Biological Process. Dot size reflects fold enrichment and colour indicates adjusted P values. **m.** Glycopeptide-level volcano plot comparing ER-G1 vs GFP facet by ER, Plasma membrane and transmembrane domain and ECM defined with g. annotation pipeline. Top-up 15 proteins are annotated.

**Supplemental Figure 6.**
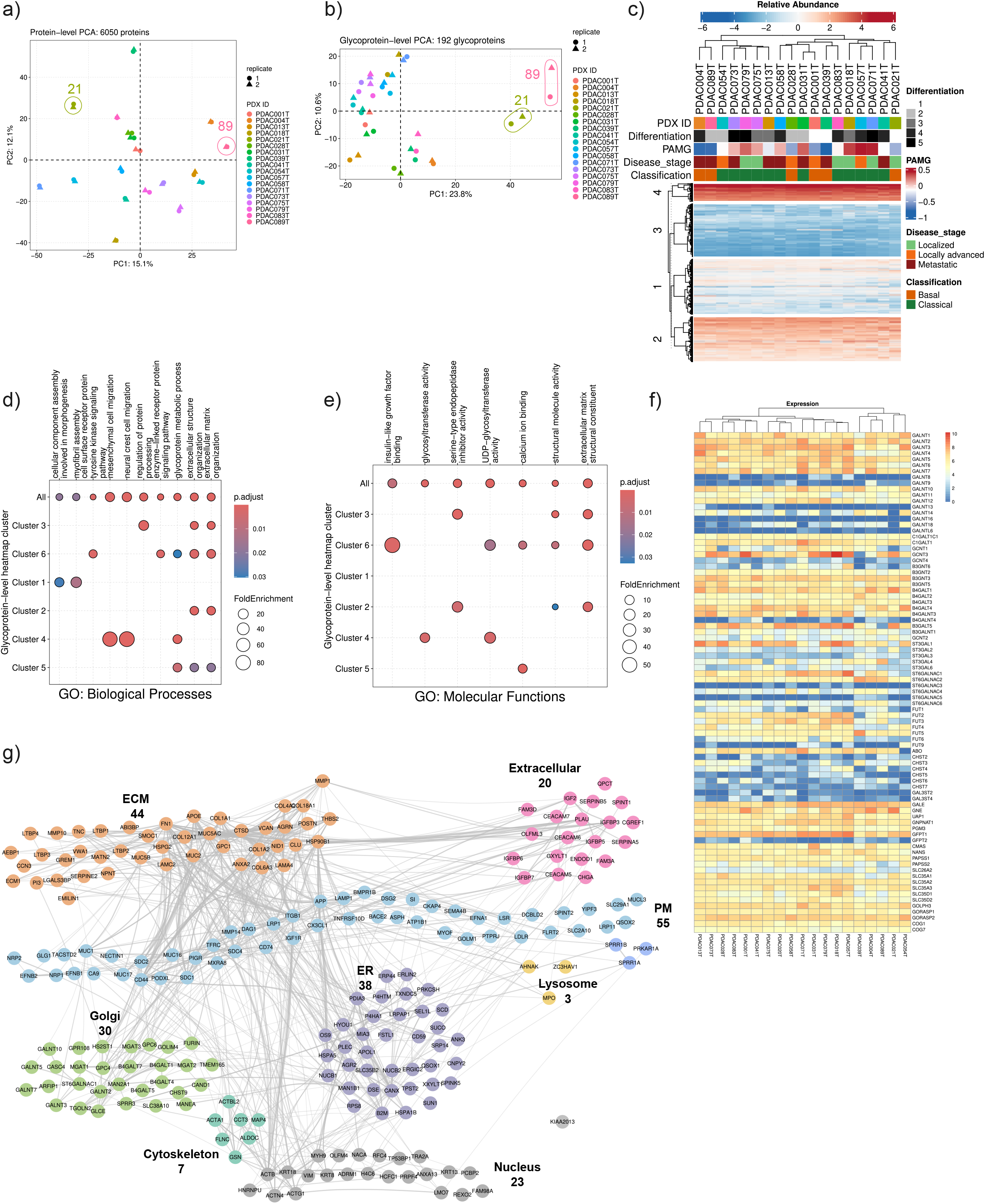
**a.** Proteome-level PCA plot for PC1 and PC2. **b.** Glyco-proteome-level PCA plot for PC1 and PC2. **c.** Proteome-level heatmap showing relative abundance comparison between PDX PDAC samples. K-means clustering was performed on protein rows. Annotations shown include differentiation levels 1-5, pancreatic adenocarcinoma molecular gradient (PAMG) from -1 (orange) to 0.5 (green), basal (blue) or classical (yellow) classification, surgery status for resection (pink) or biopsy (blue) and disease stage gradient from localised to metastatic. **d.** Proteome-level PCA plot for PC1 and PC2. **c** Glycopeptide-level heatmap. **e.** Gene ontology biological processes analysis of glycoproteins detected in PDX samples split by heatmap cluster. False discovery rate (p.adjust) shown by colour scale and gene ratio by dot size. **f.** Gene ontology molecular function analysis of glycoproteins detected in PDX samples split by heatmap cluster. False discovery rate (p.adjust) shown by colour scale and gene ratio by dot size. **g.** STRING PPI network of PDX-derived candidate proteins (n = 220, Homo sapiens), organized by subcellular compartment. Layout and parameters as in Figure 4i. **h.** Expression Heatmap.

## References

1. Siegel, R.L., Kratzer, T.B., Giaquinto, A.N., Sung, H., and Jemal, A. (2025). Cancer statistics, 2025. CA Cancer J. Clin. 75, 10–45.

2. Sherman, M.H., and Beatty, G.L. (2023). Tumor microenvironment in pancreatic cancer pathogenesis and therapeutic resistance. Annu. Rev. Pathol. 18, 123–148.

3. Ho, W.J., Jaffee, E.M., and Zheng, L. (2020). The tumour microenvironment in pancreatic cancer - clinical challenges and opportunities. Nat. Rev. Clin. Oncol. 17, 527–540.

4. Pamela Stanley, Kelley W. Moremen, Nathan E. Lewis, Naoyuki Taniguchi, and Markus Aebi. (2022). N-Glycans. In Essentials of Glycobiology 4th edition., Varki A, Cummings RD, Esko JD, et al., editors., ed. (Cold Spring Harbor Laboratory Press).

5. Springer, G.F. (1984). T and Tn, general carcinoma autoantigens. Science 224, 1198–1206.

6. Cull, J., Pink, R.C., Samuel, P., and Brooks, S.A. (2025). Myriad mechanisms: factors regulating the synthesis of aberrant mucin-type O-glycosylation found on cancer cells. Glycobiology 35. 10.1093/glycob/cwaf023.

7. Ju, T., Lanneau, G.S., Gautam, T., Wang, Y., Xia, B., Stowell, S.R., Willard, M.T., Wang, W., Xia, J.Y., Zuna, R.E., et al. (2008). Human tumor antigens Tn and sialyl Tn arise from mutations in Cosmc. Cancer Res. 68, 1636–1646.

8. Radhakrishnan, P., Dabelsteen, S., Madsen, F.B., Francavilla, C., Kopp, K.L., Steentoft, C., Vakhrushev, S.Y., Olsen, J.V., Hansen, L., Bennett, E.P., et al. (2014). Immature truncated O-glycophenotype of cancer directly induces oncogenic features. Proc. Natl. Acad. Sci. U. S. A. 111, E4066–E4075.

9. Gill, D.J., Chia, J., Senewiratne, J., and Bard, F. (2010). Regulation of O-glycosylation through Golgi-to-ER relocation of initiation enzymes. J. Cell Biol. 189, 843–858.

10. Gill, D.J., Clausen, H., and Bard, F. (2011). Location, location, location: new insights into O-GalNAc protein glycosylation. Trends Cell Biol. 21, 149–158.

11. Chia, J., Tham, K.M., Gill, D.J., Bard-Chapeau, E.A., and Bard, F.A. (2014). ERK8 is a negative regulator of O-GalNAc glycosylation and cell migration. Elife 3, e01828.

12. Chia, J., Tay, F., and Bard, F. (2019). The GalNAc-T Activation (GALA) Pathway: Drivers and markers. PLoS One 14, e0214118.

13. Chia, J., Wang, S.-C., Wee, S., Gill, D.J., Tay, F., Kannan, S., Verma, C.S., Gunaratne, J., and Bard, F.A. (2021). Src activates retrograde membrane traffic through phosphorylation of GBF1. Elife 10. 10.7554/eLife.68678.

14. Gill, D.J., Tham, K.M., Chia, J., Wang, S.C., Steentoft, C., Clausen, H., Bard-Chapeau, E.A., and Bard, F.A. (2013). Initiation of GalNAc-type O-glycosylation in the endoplasmic reticulum promotes cancer cell invasiveness. Proc. Natl. Acad. Sci. U. S. A. 110, E3152–E3161.

15. Nguyen, A.T., Chia, J., Ros, M., Hui, K.M., Saltel, F., and Bard, F. (2017). Organelle Specific O-Glycosylation Drives MMP14 Activation, Tumor Growth, and Metastasis. Cancer Cell 32, 639–653.e6.

16. Ros, M., Nguyen, A.T., Chia, J., Le Tran, S., Le Guezennec, X., McDowall, R., Vakhrushev, S., Clausen, H., Humphries, M.J., Saltel, F., et al. (2020). ER-resident oxidoreductases are glycosylated and trafficked to the cell surface to promote matrix degradation by tumour cells. Nat. Cell Biol. 22, 1371–1381.

17. Ye, Z., Mao, Y., Clausen, H., and Vakhrushev, S.Y. (2019). Glyco-DIA: a method for quantitative O-glycoproteomics with in silico-boosted glycopeptide libraries. Nat. Methods 16, 902–910.

18. Du, Z., and Lovly, C.M. (2018). Mechanisms of receptor tyrosine kinase activation in cancer. Mol. Cancer 17, 58.

19. Hamidi, H., and Ivaska, J. (2018). Every step of the way: integrins in cancer progression and metastasis. Nat. Rev. Cancer 18, 533–548.

20. Cavallaro, U., and Christofori, G. (2004). Cell adhesion and signalling by cadherins and Ig-CAMs in cancer. Nat. Rev. Cancer 4, 118–132.

21. Efthymiou, G., Lohmann, E., Montenegro, C., Kousteridou, P., Bertrand, P., Pereira, J., Carr, H.S., Ben-Sahra, I., Compare, C., Nemazanyy, I., et al. (2025). The extracellular matrix drives guanylate production and protects pancreatic cancer cells from oxaliplatin-induced DNA damage. Sci. Adv. 11, eadu2276.

22. Nicolle, R., Blum, Y., Duconseil, P., Vanbrugghe, C., Brandone, N., Poizat, F., Roques, J., Bigonnet, M., Gayet, O., Rubis, M., et al. (2020). Establishment of a pancreatic adenocarcinoma molecular gradient (PAMG) that predicts the clinical outcome of pancreatic cancer. EBioMedicine 57, 102858.

23. Mutgan, A.C., Besikcioglu, H.E., Wang, S., Friess, H., Ceyhan, G.O., and Demir, I.E. (2018). Insulin/IGF-driven cancer cell-stroma crosstalk as a novel therapeutic target in pancreatic cancer. Mol. Cancer 17, 66.

24. Katrine, T.-B.S., Vakhrushev, S.Y., Kong, Y., Steentoft, C., Nudelman, A.S., Pedersen, N.B., Wandall, H.H., Mandel, U., Bennett, E.P., Levery, S.B., et al. (2012). Probing isoform-specific functions of polypeptide GalNAc-transferases using zinc finger nuclease glycoengineered SimpleCells. Proceedings of the National Academy of Sciences 109, 9893–9898.

25. Steentoft, C., Vakhrushev, S.Y., Joshi, H.J., Kong, Y., Vester-Christensen, M.B., Katrine, T., Schjoldager, B.G., Lavrsen, K., Dabelsteen, S., Pedersen, N.B., et al. (2013). Precision mapping of the human O-GalNAc glycoproteome through SimpleCell technology. EMBO J. 32, 1478–1488.

26. Cheng, J.J., Matsumoto, Y., Dombek, G.E., Stackhouse, K.A., Ore, A.S., Glickman, J.N., Heimburg-Molinaro, J., and Cummings, R.D. (2025). Differential expression of CD175 and CA19-9 in pancreatic adenocarcinoma. Sci. Rep. 15, 4177.

27. Ju, T., Aryal, R.P., Kudelka, M.R., Wang, Y., and Cummings, R.D. (2014). The Cosmc connection to the Tn antigen in cancer. Cancer Biomark. 14, 63–81.

28. Thomas, D., Sagar, S., Caffrey, T., Grandgenett, P.M., and Radhakrishnan, P. (2019). Truncated O-glycans promote epithelial-to-mesenchymal transition and stemness properties of pancreatic cancer cells. J. Cell. Mol. Med. 23, 6885–6896.

29. Chia, J., Goh, G., and Bard, F. (2016). Short O-GalNAc glycans: regulation and role in tumor development and clinical perspectives. Biochim. Biophys. Acta 1860, 1623–1639.

30. Hodgson, K., Orozco-Moreno, M., Peng, Z., Blencoe, L., Sharp, M.J., Smith, E., Grimsley, G., Elliott, D.J., Beatson, R., Drake, R.R., et al. (2026). The glycobiology of prostate cancer: an update. Oncogene. 10.1038/s41388-026-03858-x.

31. Mohamed Abd-El-Halim, Y., El Kaoutari, A., Silvy, F., Rubis, M., Bigonnet, M., Roques, J., Cros, J., Nicolle, R., Iovanna, J., Dusetti, N., et al. (2021). A glycosyltransferase gene signature to detect pancreatic ductal adenocarcinoma patients with poor prognosis. EBioMedicine 71, 103541.

32. Rodriguez, E., Boelaars, K., Brown, K., Madunić, K., van Ee, T., Dijk, F., Verheij, J., Li, R.J.E., Schetters, S.T.T., Meijer, L.L., et al. (2022). Analysis of the glyco-code in pancreatic ductal adenocarcinoma identifies glycan-mediated immune regulatory circuits. Commun. Biol. 5, 41.

33. Hodgson, K., Orozco-Moreno, M., Scott, E., Garnham, R., Livermore, K., Thomas, H., Zhou, Y., He, J., Bermudez, A., Garcia Marques, F.J., et al. (2023). The role of GCNT1 mediated O-glycosylation in aggressive prostate cancer. Sci. Rep. 13, 17031.

34. Joshi, H.J., Narimatsu, Y., Schjoldager, K.T., Tytgat, H.L.P., Aebi, M., Clausen, H., and Halim, A. (2018). SnapShot: O-glycosylation pathways across kingdoms. Cell 172, 632–632.e2.

35. Schjoldager, K.T., Narimatsu, Y., Joshi, H.J., and Clausen, H. (2020). Global view of human protein glycosylation pathways and functions. Nat. Rev. Mol. Cell Biol. 21, 729–749.

36. Borrok, M.J., Jung, S.T., Kang, T.H., Monzingo, A.F., and Georgiou, G. (2012). Revisiting the role of glycosylation in the structure of human IgG Fc. ACS Chem. Biol. 7, 1596–1602.

37. Spiteri, V.A., Doutch, J., Rambo, R.P., Gor, J., Dalby, P.A., and Perkins, S.J. (2021). Solution structure of deglycosylated human IgG1 shows the role of CH2 glycans in its conformation. Biophys. J. 120, 1814–1834.

38. Arnold, J.N., Wormald, M.R., Sim, R.B., Rudd, P.M., and Dwek, R.A. (2007). The impact of glycosylation on the biological function and structure of human immunoglobulins. Annu. Rev. Immunol. 25, 21–50.

39. Nath, S., and Mukherjee, P. (2014). MUC1: a multifaceted oncoprotein with a key role in cancer progression. Trends Mol. Med. 20, 332–342.

40. Dolgin, E. (2023). Disrupting protein folding to tackle cancer. Nature. 10.1038/d41586-023-01649-y.

41. Youssef, K.K., Narwade, N., Arcas, A., Marquez-Galera, A., Jiménez-Castaño, R., Lopez-Blau, C., Fazilaty, H., García-Gutierrez, D., Cano, A., Galcerán, J., et al. (2024). Two distinct epithelial-to-mesenchymal transition programs control invasion and inflammation in segregated tumor cell populations. Nat. Cancer 5, 1660–1680.

42. Jakobsen, S.T., and Siersbæk, R. (2025). Transcriptional regulation by MYC: an emerging new model. Oncogene 44, 1–7.

43. Phan, L.M., and Rezaeian, A.-H. (2021). ATM: Main features, signaling pathways, and its diverse roles in DNA damage response, tumor suppression, and cancer development. Genes (Basel) 12, 845.

44. Guillaumond, F., Bidaut, G., Ouaissi, M., Servais, S., Gouirand, V., Olivares, O., Lac, S., Borge, L., Roques, J., Gayet, O., et al. (2015). Cholesterol uptake disruption, in association with chemotherapy, is a promising combined metabolic therapy for pancreatic adenocarcinoma. Proc. Natl. Acad. Sci. U. S. A. 112, 2473–2478.

45. Dobin, A., Davis, C.A., Schlesinger, F., Drenkow, J., Zaleski, C., Jha, S., Batut, P., Chaisson, M., and Gingeras, T.R. (2013). STAR: ultrafast universal RNA-seq aligner. Bioinformatics 29, 15–21.

46. Liao, Y., Smyth, G.K., and Shi, W. (2014). featureCounts: an efficient general purpose program for assigning sequence reads to genomic features. Bioinformatics 30, 923–930.

47. Bullard, J.H., Purdom, E., Hansen, K.D., and Dudoit, S. (2010). Evaluation of statistical methods for normalization and differential expression in mRNA-Seq experiments. BMC Bioinformatics 11, 94.

